# Multi-trophic risk from human superpredators may alter predator-prey coexistence and population dynamics

**DOI:** 10.64898/2026.06.12.731855

**Authors:** Shawn Dsouza

**Affiliations:** Center for Ecological Sciences, Indian Institute of Science, Bengaluru, India

**Keywords:** agent-based model, non-consumptive effects, risk-disturbance hypothesis, trophic downgrading, trophic cascade

## Abstract

Humans are efficient and deadly predators, yet they may also interact with wildlife in non-lethal ways. This study explores how interactions with lethal and non-lethal human “superpredators” alter predator-prey dynamics using an agent-based modelling approach. Our model incorporates both the consumptive (lethal) and non-consumptive (behavioural) effects of humans, as well as of predators on prey. We explored how the replacement of apex predators by humans affects mesopredator–prey dynamics, with particular emphasis on trophic targeting and differences between lethal and non-lethal interactions. We found that human superpredators have a greater effect on model outcomes than apex predators. When superpredators consume mesopredators alone or with prey, the probability of mesopredator-prey coexistence increases to a greater extent than when apex predators consume mesopredators. In contrast, superpredators consuming only prey slightly increases overall extinction risks and reduces coexistence. Non-lethal superpredators, despite eliciting anti-predator responses in mesopredators and prey, had a negligible effect on population dynamics. Our findings demonstrate that human superpredators may functionally replace apex predators when they are lethal. However, non-lethal interactions with humans may not be as ecologically significant as lethal interactions, even when humans induce anti-predator responses.

## Introduction

Predators and prey are locked in a continuous behavioural response race, where predators strive to capture more prey, and prey attempt to avoid capture (Sih, 1984). The players in this ecological game influence each other’s spatial distributions and population dynamics by altering their behaviours over time. Humans may also act as predators in many ecosystems, but they are significantly more deadly than other predators. Thus, human predators have been referred to as “superpredators” (Darimont et al., 2015, 2023). A recent meta-analysis suggests that wild animals significantly alter their behaviour in response to lethal human superpredators and to a lesser extent to non-lethal humans in their environment (Dsouza et al., 2025). However, the effect of human superpredators on predator-prey dynamics remains understudied and lacks comprehensive theoretical exploration (Smith et al., 2024)

The dynamics of the predator-prey game are governed by the energy requirements of predators and prey, the risk of predation experienced by prey, as well as the anti-predator strategies employed by prey (Brown et al., 1999; Brown & Kotler, 2004; Lima & Dill, 1990). The risk allocation hypothesis posits that prey must balance their energy and mating needs with the risk of predation (Brown et al., 1999; Lima & Bednekoff, 1999; e.g., Heithaus & Dill, 2002). To optimise this trade-off, prey adopt a variety of morphological, physiological, and behavioural strategies (Werner & Peacor, 2003). Accordingly, theoretical work has sought to formalise how prey integrate information about predation risk into behavioural decision-making.

Lima and Dill (1990) explored behavioural decisions under predation risk, presenting a mechanistic, hierarchical model that outlines branching outcomes when prey encounter predators. This model provides a probabilistic framework for understanding how prey weigh their decisions in such scenarios and extends to foraging strategies in patchy environments (see Figure 1 in Lima and Dill, 1990). Behaviourally responsive prey can leverage information about predators in their environment to increase their vigilance, enhancing their ability to detect predators. However, this comes at the cost of reduced foraging time (Brown & Kotler, 2004). Alternatively, prey may simply relocate to safer habitats to avoid predators (Luttbeg & Schmitz, 2000). While the cost of such anti-predator strategies may reduce prey survival and fecundity, thereby influencing population dynamics (see Sheriff et al., 2020 for critique), the life-dinner principle suggests that the fitness cost of anti-predator strategies is negligible compared to the absolute cost of death from predation (Dawkins & Krebs, 1997).

**Figure 1:**
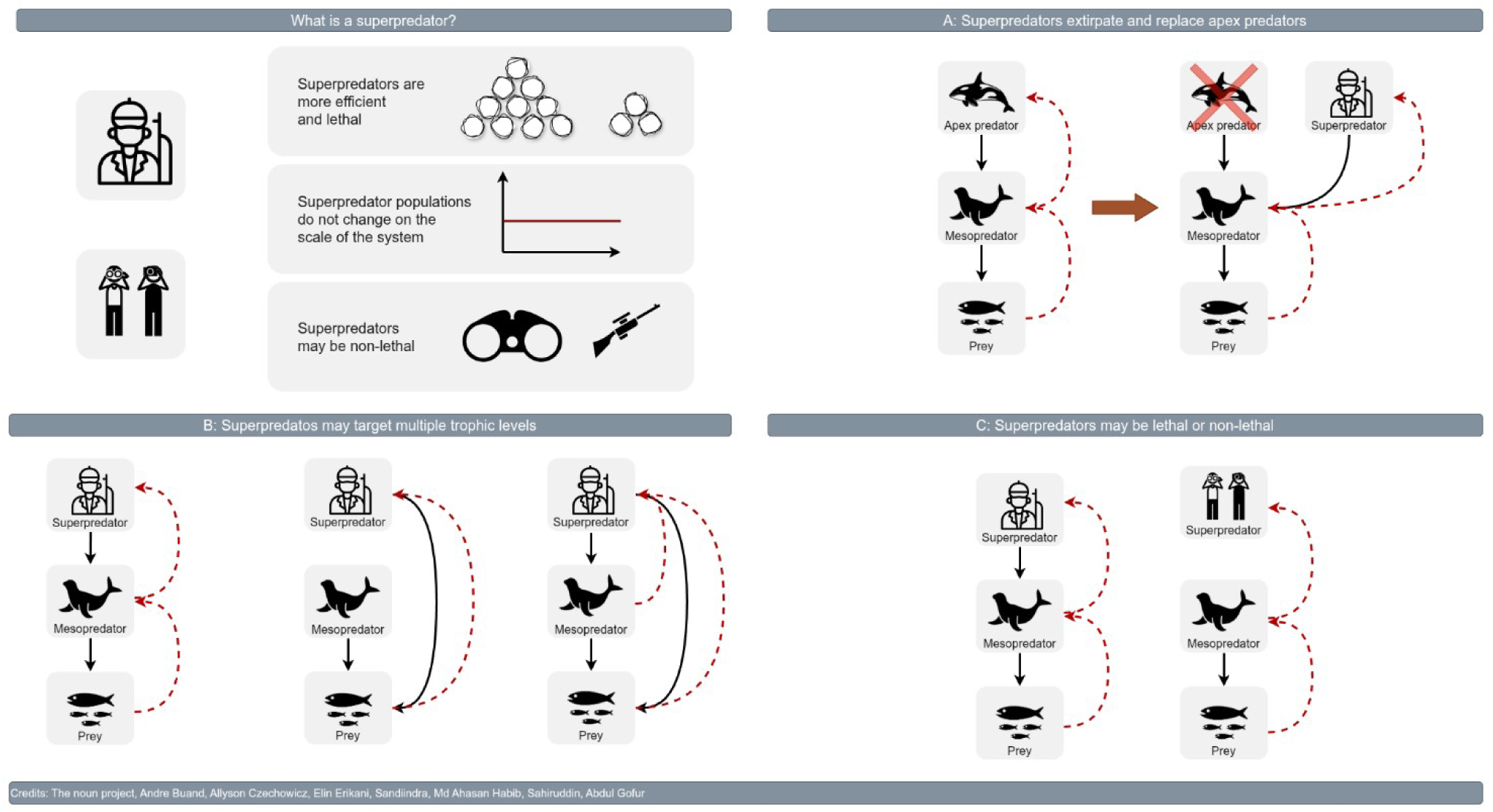
Defining superpredators (top left) and the possible modalities through which human superpredators may interact with predator – prey systems. A) Extirpating and replacing apex predators, targetting mesopredators B) Targeting one or both trophic levels C) Benign interactions with wild animals such as ecotourism. Black arrows indicate the direction of consumption or harvesting. Red dashed arrows indicate the direction of anti-predator responses. (Icon credits: the noun project, Andre Buand, Allyson Czechowicz, Elin Erikani, Sandiindra, Credits: Md Ahasan Habib, Sahiruddin, Abdul Gofur).

Predators, too, can adapt their behaviour to increase prey encounters. For example, predators targeting sessile or slow-moving prey may adopt a random or area-restricted movement strategy, which has been shown to be evolutionarily stable (Mitchell & Lima, 2002). Such search behaviour may maximise encounter rates when prey distribution is patchy or unpredictable. However, when prey can behaviourally avoid predators in real time, predators may use more sophisticated hunting tactics. Predators may use spatial information such as successful hunting sites from past interactions, focus on areas with higher prey densities, or use environmental cues such as scent and auditory cues to increase their probability of locating prey hotspots. In this context, the availability of information to both predators and prey can significantly alter the dynamics and stability of the system (Luttbeg & Schmitz, 2000).

Human tool use, cooperation, and intelligence render them far more efficient than other predators (Darimont et al., 2015; Treves & Naughton-Treves, 1999). Human exploitation is dramatically more extensive compared to non-human predators, affecting roughly a third of all vertebrate species, often for non-food reasons such as the pet trade, medicine, or ornamental uses. Unlike other predators, human population dynamics are decoupled from the species and systems they interact with, at least on the ecological timescale of these interactions. This absence of feedback tends to facilitate the overexploitation of targeted species (Darimont et al., 2023).

Humans tend to disproportionately target predator species for their higher economic value (Pauly et al., 1998). This has led to the extirpation of these predator species in some environments due to overexploitation (Estes et al., 2011; Jackson et al., 2001). Such trophic downgrading has downstream effects for ecosystems. For example, the removal of sea otters (*Enhydra lutris*) released sea urchin populations from top – down control. The resulting population explosion of sea urchins and increased herbivory led to the eventual collapse of kelp forests along the Alaska coast (Estes & Duggins, 1995). In other systems, humans may exacerbate existing predation risks (Smith et al., 2024; Dsouza et al., 2025). For example, human hunters coexist with wolves (*Canis lupus*) in North America and Europe and share a common ungulate prey base such as elk (*Cervus elaphus*), roe deer (*Capreolus capreolus*) and wild boar (*Sus scrofa*). During hunting seasons these prey populations experience added risk from human hunters in addition to wolves (Wenting et al., 2024).

Humans can also interact with ecosystems in non-lethal ways, such as through tourism, hiking, or even walking in parks (e.g., Fernandez-Juricic et al., 2003). The risk-disturbance hypothesis suggests that some animals respond to such interactions strongly, in ways that are akin to how they respond to natural predators (Frid & Dill, 2002). For example, yellow-bellied marmots (*Marmota flaviventris*) in North America reduced foraging and increased vigilance significantly in response to benign humans in their environment (Uchida & Blumstein, 2021). However, animal responses to non-lethal humans in their environment are smaller than their responses to lethal humans (Dsouza et al., 2025). Humans may also act as shields deterring predators and reducing risk for prey species (Gaynor et al., 2025). This mismatch between the actual risk posed by these interactions and the animals’ responses can impose undue fitness costs, potentially affecting the overall population dynamics of prey species (Smith et al., 2021).

Our goal is to understand how interactions with human “superpredators” alter predator-prey dynamics in different contexts (Figure 1). We use an agent-based model with prey, mesopredators, and apex predators that are either natural or exhibit human superpredator characteristics. These agents can sense and respond to each other. Through simulation experiments, we evaluate the population trajectories of these agents and the system’s stable states in the following scenarios (Table 1):

1. a) A specialist apex predator consumes only mesopredators, which in turn target prey species or (b) a generalist apex predator targets both prey and mesopredators. The apex predator is not fully efficient (15% success rate), and its population dynamics are tied to the population dynamics of its chosen target, i.e., mesopredators.
2. The apex predator is replaced by a superpredator that is 100% efficient and whose population remains constant over time. The superpredators that target (a) mesopredators, (b) prey, or (c) both.
3. Non-lethal superpredators which do not consume any agents but elicit anti-predator responses.

**Table 1:** Description of scenarios simulated in the current study.

| Scenario | Description |
| --- | --- |
| Baseline | Mesopredator consumes prey |
| 1 | Apex predator consumes mesopredator |
| 2 | Apex predator consumes both prey and mesopredator |
| 3 | Superpredator consumes prey |
| 4 | Superpredator consumes mesopredator |
| 5 | Superpredator consumes both prey and mesopredator |
| 6 | Prey respond to non-lethal superpredator |
| 7 | Mesopredator respond to non-lethal superpredator |
| 8 | Both prey and mesopredator respond to non-lethal superpredator |

## Methods

### Model description

The model description follows the ODD (Overview, Design Concepts, Details) protocol for describing individual- and agent-based models (Grimm et al., 2006, updated 2020). We implemented the model using the Mesa agent-based modelling framework in python 3.12.5 (Masad & Kazil, 2015). The code for our implementation is available at https://doi.org/10.5281/zenodo.20666048.

### Purpose and patterns

The purpose of this model is to explore how the dynamics of a simple Leslie-Gower-type predator-prey system are altered by interactions with a superpredator. The superpredator is defined as a species of agent that may target predator agents, prey agents, or both, and can be either lethal or non-lethal depending on the scenario being tested. Lethal superpredators remove other agents (mesopredator, prey or both) that they encounter and elicit anti-predator responses depending on the scenario being simulated. In contrast, the non-lethal superpredators only elicit anti-predator responses from the agents they encounter. We focus on population-level outcomes, specifically: when only prey survive to the end (prey only), when mesopredators consume all prey (extinction), and when both mesopredators and prey survive to the end (coexistence). Additionally, we analyse the steady-state dynamics, such as population cycles, under coexistence regimes. This allows us to understand how superpredators influence the stability and persistence of predator-prey systems.

### Entities, state variables and scales

The model consists of four types of agents: prey, mesopredators, apex predators, and superpredators. These agents exist on a discrete square lattice (*L2*) of 100 × 100 cells with periodic boundary conditions, which we refer to as the environment (See Singh et al. 2024 for a recent review of spatially explicit predator – prey models). The spatial scale of the model is general but for clarity we use the example of grassland or a seagrass meadow environment. In such a case, each cell in the environment corresponds to an area of approximately 500 m², such that the full lattice covers about 5 km². However, the periodic boundary conditions make the space effectively much larger. The model is implemented in discrete time, with each time step corresponding roughly to one week. Each simulation run spans 1000 time steps, equivalent to approximately 19 years. There is no a priori limit on the number of agents that can occupy a given cell.

All agents are characterized by three core attributes: a unique identification number, their type (prey, mesopredator, apex predator, or superpredator), and their location on the lattice (defined by *x* and *y* coordinates). In addition, mesopredator and apex predator agents possess an energy state variable (*E*), represented as a non-negative integer which varies over time with an upper bound (*Emax*).

### Process overview and scheduling

At each time step, the agents in the model are activated in random order, and the model is updated asynchronously. Once activated, each agent performs a series of actions in a predetermined sequence based on its type and the local environment, which includes the eight adjacent cells. The actions, executed in the following order, include moving, hunting (for predator-type agents), avoiding predation (for prey and mesopredator agents), eating, reproducing, dying (if conditions for death are met). The specific set of actions for each type of agent is detailed in Figure 2.

**Figure 2:**
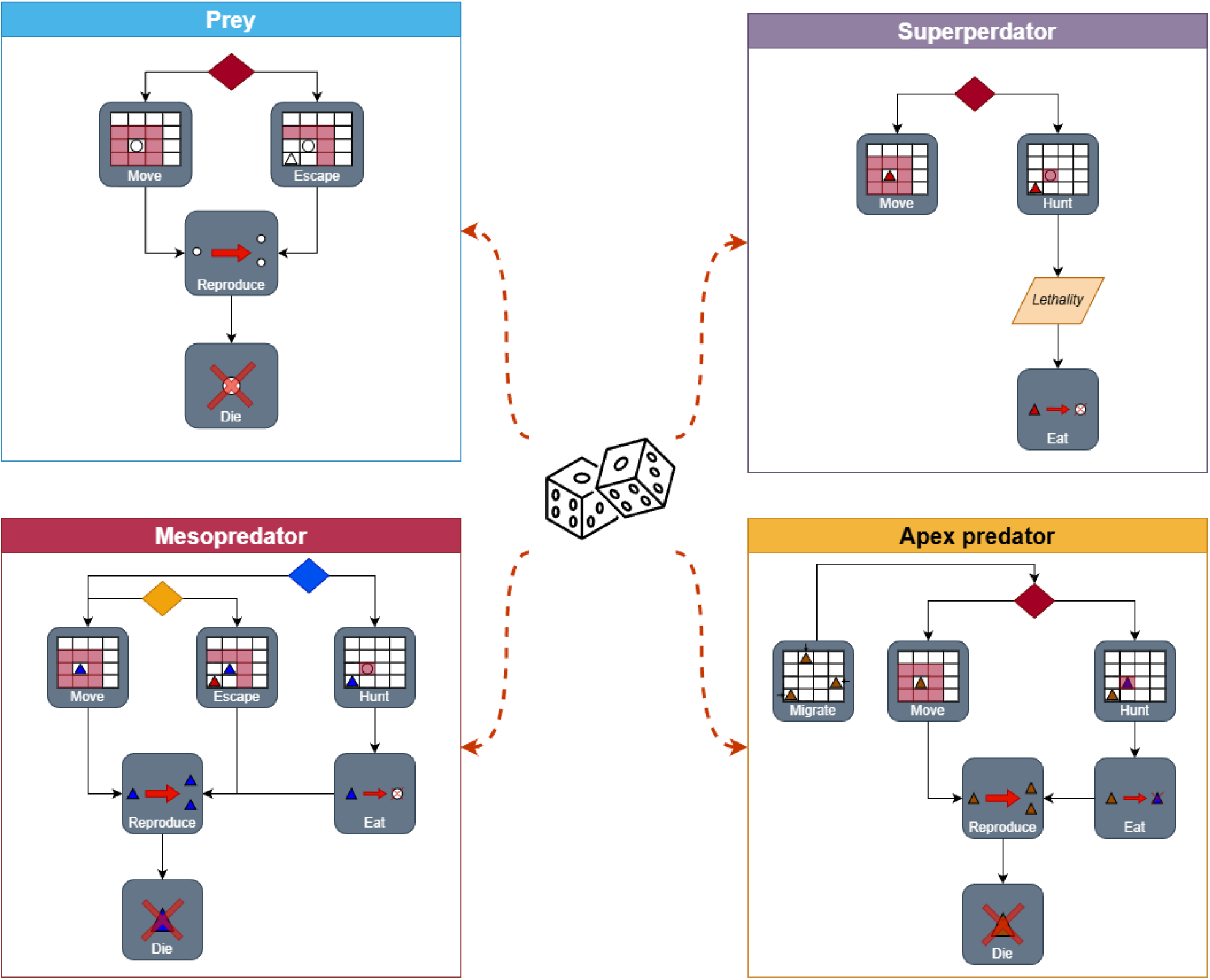
Description of model processes and agent sub-models. Agents are activated randomly and then follow a set order of actions conditional on the presence of other agents in the immediate neighborhood (colored diamonds) and traits (target and lethality for superpredators).

### Design concepts and principles

The core principle governing agent behaviour and interactions is the foraging-safety trade-off: agents must acquire energy to survive and reproduce while simultaneously avoiding predation (Brown et al., 1999; Lima & Dill, 1990). Prey agents implement behavioural anti-predator strategies, such as avoiding predators in space and time, which impose associated costs. Similarly, predator agents adopt sensory and movement strategies to increase their encounters with prey (Mitchell & Lima, 2002).

The emergent population dynamics of the model are designed to approximate a Leslie-Gower-like system (Colon et al., 2015). In this framework, prey populations grow logistically in the absence of predators, while predator populations decline and go extinct in the absence of prey. Dynamics arise from the interactions between prey and predators, primarily through avoidance behaviour by prey and attraction behaviour by predators. The model does not include any adaptive elements in agent behaviour, which means that agents do not learn or change their strategies over time. Additionally, agents do not have specific objectives and cannot predict or anticipate the actions of other agents or changes in the environment. Instead, agents operate based on instantaneous sensory information, which includes the type, number, and position of other agents within their Moore neighbourhood, consisting of the eight adjacent cells on the lattice.

Interactions between agents are structured based on their roles. Prey agents do not interact with each other but can avoid predator agents in their vicinity to reduce predation risk. Predator agents, on the other hand, can hunt and consume prey agents. Stochasticity is a key feature of the model, implemented through random processes such as birth and death events, agent movement on the lattice, and decision-making related to hunting and avoidance. This randomness introduces variability in agent behaviour and population dynamics, reflecting the inherent unpredictability of ecological systems.

At each time step, the position of every agent on the lattice is tracked, along with the total count of all types of agents present. These data allow for the analysis of population trajectories, steady-state dynamics, and the impact of superpredator interactions on the system. Collectively, these design concepts and principles ensure that the model captures the essential features of predator-prey dynamics while remaining computationally tractable for simulation experiments.

### Initialization

At the start of each simulation, agents are placed at random locations on the lattice. The initial population sizes are as follows: 500 mesopredators, 500 prey, 500 apex predators, and 100 superpredators. These initial values were determined through preliminary simulation tests to ensure the possibility of coexistence among the agent types and to generate the most ecologically interesting dynamics (see Appendix A for details).

### Prey rules

1. Move: In the absence of predators in the Moore neighbourhood (i.e., the eight neighboring cells, figure 2), prey move in a random walk across the lattice, regardless of whether cells are occupied other prey agents. Prey move a fixed number of spatial steps (*qprey*) each time step, choosing a random direction for each step.
2. Escape: When mesopredators or superpredators are present in their Moore neighborhood, prey attempt to move away from them. If multiple predators are present, prey randomly select one predator from their vicinity and move away from it. “Away” is defined as moving into the half of the Moore neighborhood opposite the predator. Each avoidance action incurs a cost (*c*) to the prey’s reproductive potential.
3. Reproduce: Prey agents can reproduce with an intrinsic probability (*bprey*), provided the number of prey in their immediate vicinity is below the saturation density (*S*). The probability of reproduction decreases with the cost of avoidance (*b′prey* − *c*) each time avoidance is implemented. If prey do not avoid predators, their reproductive probability recovers (*b′prey*+*c*) at each time step, up to the maximum intrinsic value (*bprey*).
4. Die: At each time step, prey agents have a constant intrinsic probability (*dprey*) of dying independent of predation and being removed from the lattice.

### Mesopredator rules

1. Move: In the absence of prey, mesopredators move in a random walk across the lattice, regardless of whether cells are occupied. Mesopredators move a fixed number of spatial steps (*qpredator*) each time step.
2. Hunt: When prey are present in their vicinity, mesopredators move toward them. If multiple prey are present, the mesopredator selects one randomly following a uniform probability and moves toward it.
3. Escape: When apex predators or superpredators are present in the immediate vicinity, mesopredators attempt to move away from them, using a similar strategy as prey to avoid predation. As above, each avoidance incurs a cost (*cmesopredator*)
4. Eat: When a mesopredator and prey occupy the same cell, the mesopredator may attempt to consume the prey, with some fixed probability of success (*l*) which can be viewed as the lethality of the mesopredator. If multiple prey are present, the mesopredator selects one prey individual uniformly at random for a predation attempt. Successful consumption increases the mesopredator’s energy (*E*), up to a maximum (*Emax*).
5. Reproduce: Mesopredator agents reproduce with an intrinsic probability (*bmesopredator*) for each unit of energy they possess. Similar to prey agents, the probability of reproduction for mesopredators decreases with the cost of avoiding apex predators (*b′mesopredator* − *c*) at each time step when avoidance is implemented. If mesopredators do not avoid apex predators, their reproductive probability recovers (*b′mesopredator*+*c*) at each time step, up to the maximum intrinsic value (*bmesopredator*).
6. Die: At each time step, mesopredators have an intrinsic probability of dying (*dmesopredator*) and being removed from the lattice.

### Apex predator rules

1. Move: Apex predators move in a random walk across the lattice in the absence of mesopredators for a fixed number of spatial steps (*qapex*) per time step.
2. Hunt: Apex predators target mesopredators and/or prey in their vicinity, moving toward them if present.
3. Eat: When an apex predator and mesopredator and/or prey occupy the same cell, the apex predator may consume the mesopredator and/or prey, thereby gaining energy. The lethality level (*lapex*) of apex predators determines the success of each encounter. Apex predator gain one unit of energy for every mesopredator they consume up to a maximum of *Emax*. We assume an apex predator success rate of 15% based on empirical estimates (e.g., Standers et al., 1978, MacNulty et al. 2011, Hayward et al. 2014) and our testing (Appendix A).
4. Reproduce: Apex predator agents reproduce with an intrinsic probability (*bapex*) for each unit of energy they possess.
5. Die: Apex predators have an intrinsic probability of dying (*dapex predator*)and being removed from the lattice at each time step.
6. Migrate: We implemented a rule to ensure apex predators were present throughout the simulation in order to facilitate comparisons between apex predators and superpredators (Figure 1). When the number of apex predators on the lattice falls below a threshold value (N = 250), a small number of apex predators (N = 10) are introduced at the edges of the environment to avoid the extinction of apex predator agents.

### Superpredator rules

1. Move: Superpredators move in a random walk across the lattice or may actively seek out their targets (mesopredators or prey) depending on the scenario.
2. Hunt: Superpredators target mesopredators, prey, or both, depending on the scenario being tested and move towards them in their vicinity.
3. Eat: When superpredators are designated as lethal, they consume their targets (mesopredators or prey) with 100% success whenever they occupy the same cell. When superpredators are non-lethal, they do not consume their targets, but their presence still influences the behavior of other agents in the system through the target’s avoidance and escape rules.

### Simulation experiments

For each scenario described previously (Figure 1), we conducted 25 simulations, varying the birth probabilities of prey and mesopredators between 0.1 and 1 in increments of 1/50. We conducted extensive sensitivity testing of other parameter such as the initial number of agents, size of the lattice, lethality of mesopredator and apex predator agents and the handling limits of apex predator and mesopredator agents. Sensitivity testing was carried out by varying each paramter in fixed steps indepently in separate simulation (Appendix A). The final parameter values (Table 2) were selected on the basis of preliminary tests that identified the conditions producing the most ecologically interesting dynamics, defined as those in which mesopredators and prey were most likely to coexist. We used the scenario with only prey and mesopredator agents as a control, comparing changes in our outcomes of interest across scenarios. To focus the analysis on dynamics near the model’s steady state, we discarded the first 400 time steps of each simulation. The final state of each simulation was classified into one of three categories: Prey Only (only prey remained), Coexistence (both prey and mesopredators persisted), or Extinction (no agents remain) (Colon et al., 2015). We report the proportion of realisations(*P*) in which each state was observed for a given combination of prey and mesopredator birth probabilities. For simulations where coexistence was observed, we analysed the power spectrum of population time series to identify cycles and dominant frequencies. We compared the parameter space (*bmesopredator* vs. *bprey*) required for each state and the marginal probability of each state. Additionally, for simulations with coexistence, we evaluated the power spectra of population dynamics to assess differences in the dominant period of population fluctuations across scenarios.

**Table 2:**
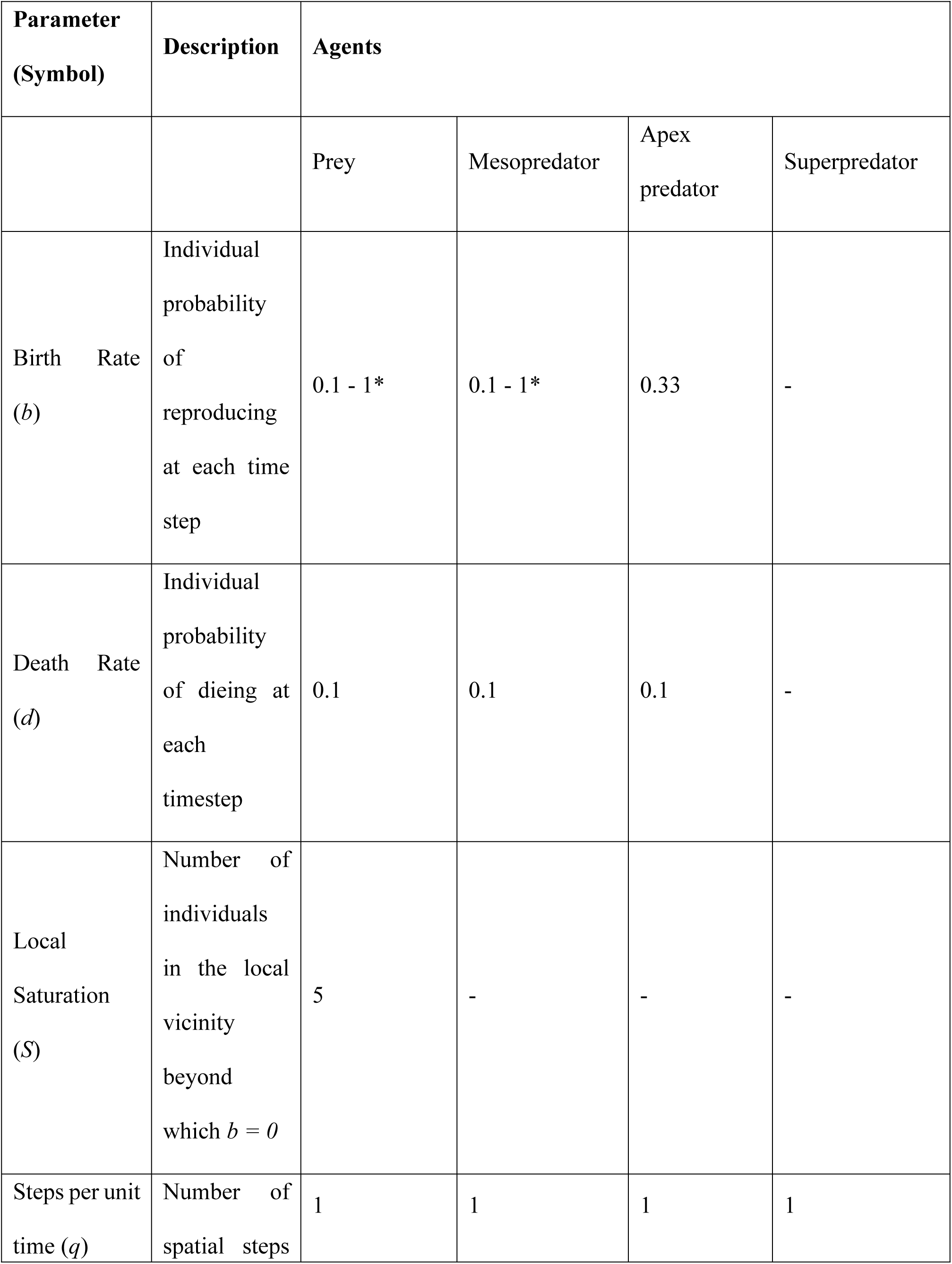
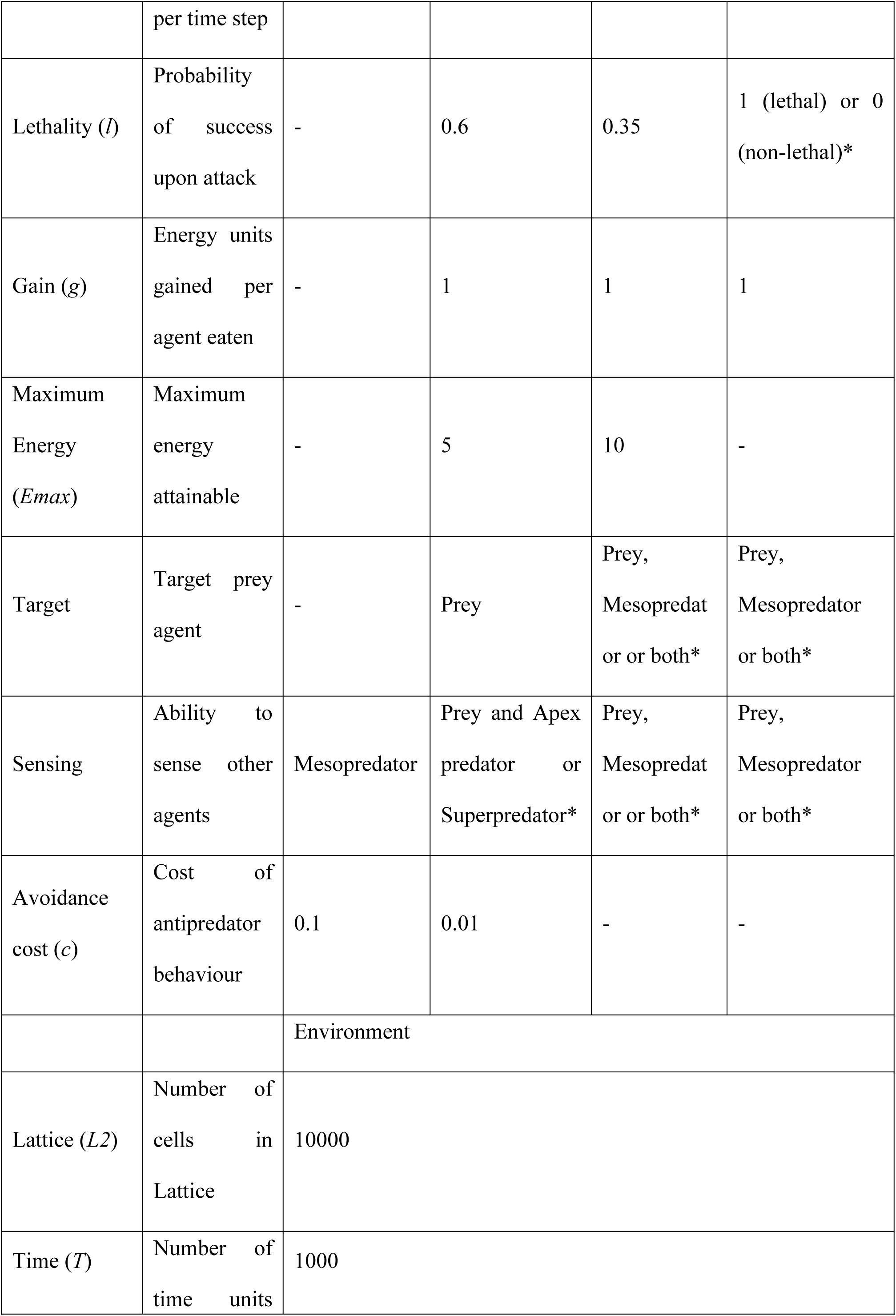
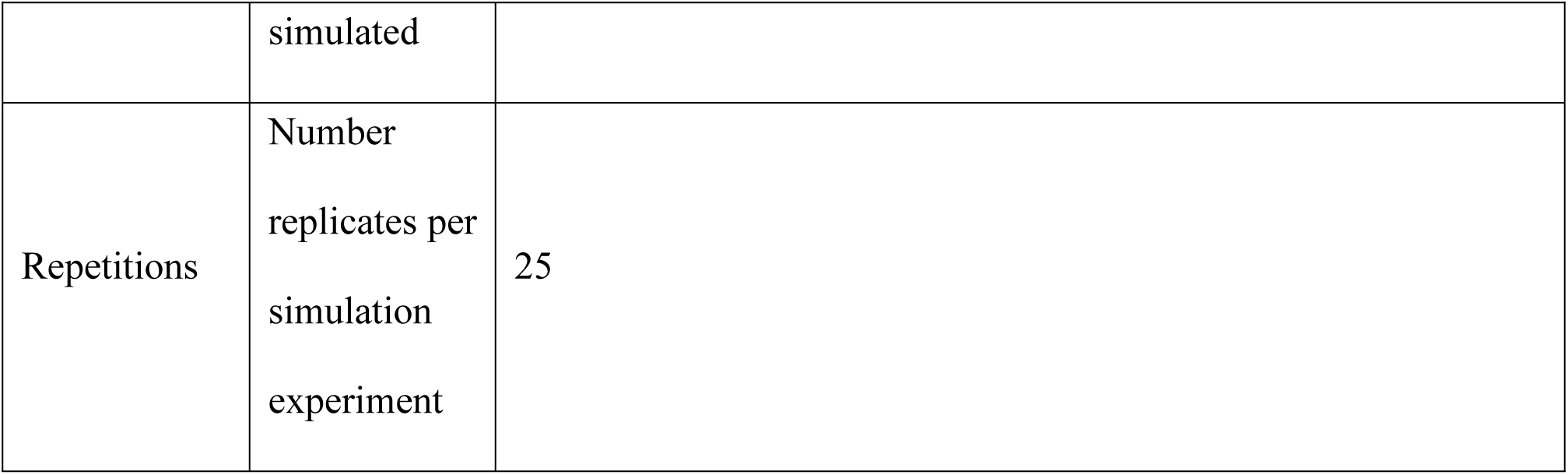
Description of default parameters of agents and environment used for simulations experiments. Values marked ‘*’ are varied during simulation experiments. Values marked ‘-‘ do not apply to that agent type.

| Parameter<br>(Symbol) | Description | Agents |  |  |  |
| --- | --- | --- | --- | --- | --- |
|  |  | Prey | Mesopredator | Apex<br>predator | Superpredator |
| Birth Rate<br>(*b*) | Individual probability of reproducing at each time step | 0.1 - 1* | 0.1 - 1* | 0.33 | - |
| Death Rate<br>(*d*) | Individual probability of dieing at each timestep | 0.1 | 0.1 | 0.1 | - |
| Local Saturation<br>(*S*) | Number of individuals in the local vicinity beyond which $b = 0$ | 5 | - | - | - |
| Steps per unit time (*q*) | Number of spatial steps per time step | 1 | 1 | 1 | 1 |
| Lethality (*l*) | Probability of success upon attack | - | 0.6 | 0.35 | 1 (lethal) or 0 (non-lethal)* |
| Gain (*g*) | Energy units gained per agent eaten | - | 1 | 1 | 1 |
| Maximum Energy (*E<sub>max</sub>*) | Maximum energy attainable | - | 5 | 10 | - |
| Target | Target prey agent | - | Prey | Prey, Mesopredator or both* | Prey, Mesopredator or both* |
| Sensing | Ability to sense other agents | Mesopredator | Prey and Apex predator or Superpredator* | Prey, Mesopredator or both* | Prey, Mesopredator or both* |
| Avoidance cost (*c*) | Cost of antipredator behaviour | 0.1 | 0.01 | - | - |
|  |  | Environment |  |  |  |
| Lattice (*L2*) | Number of cells in Lattice | 10000 |  |  |  |
| Time (*T*) | Number of time units | 1000 |  |  |  |

|  |  |  |
| --- | --- | --- |
|  | simulated |  |
| Repetitions | Number replicates per simulation experiment | 25 |

## Results

In the baseline scenario with only mesopredators consuming prey, coexistence is the most likely state (P = 0.779), followed by the prey only (P=0.132) and the extinction state (P=0.089). Coexistence occurs over a broader range of predator and prey birth rates compared to the prey only and extinction states (Figure 3, Appendix B: Table B2). When coexistence is observed, populations of both predators and prey exhibit oscillations, with a dominant period of 107 weeks for prey and 101 weeks for predators (Appendix C).

**Figure 3:**
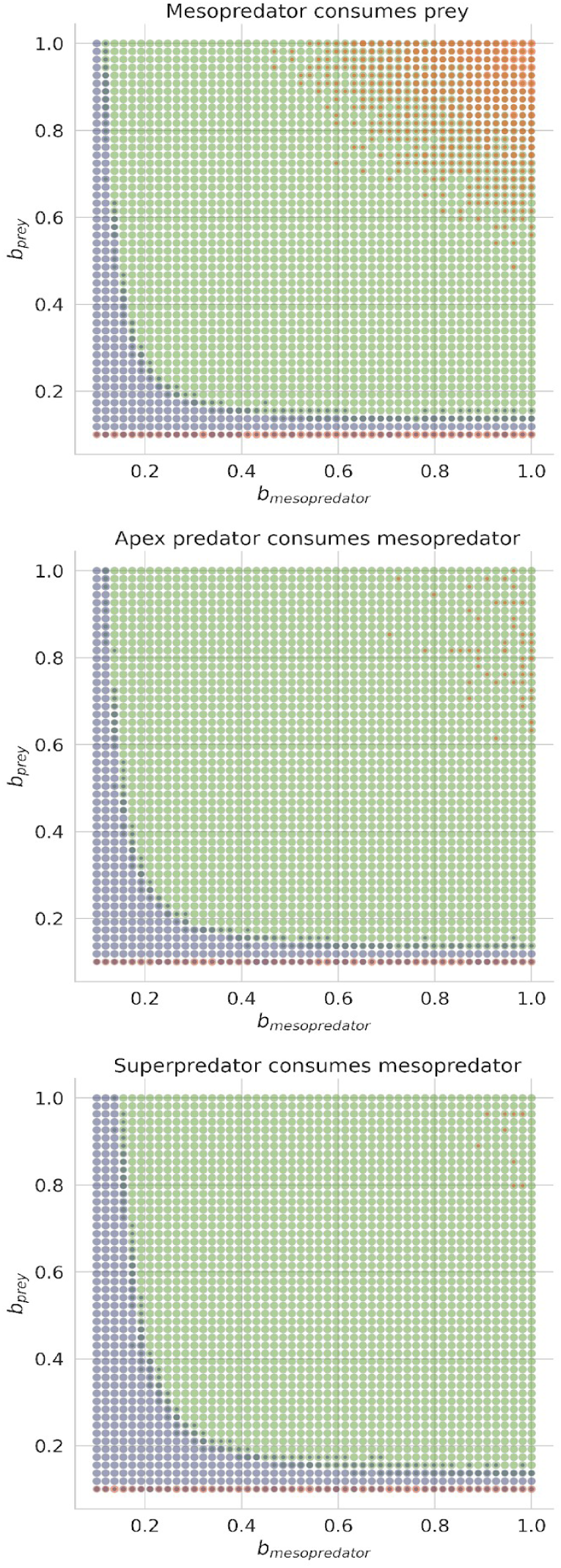
Outcomes of 25 repeated simulations across varying prey birth rates (bprey) and mesopredator birth rates (bmesopredator) under three scenarios: (1) mesopredators consuming prey, (2) apex predators consuming mesopredators, and (3) superpredators consuming mesopredators. Colours indicate the proportion of realisations resulting in Prey Only (blue), Coexistence (green), and Extinction (orange) across the parameter space.

### Adding specialist and generalist apex predators

In the absence of migration, specialist apex predators go extinct across all parameter values tested (see Appendix A for model validation results). Accordingly, our analyses focus on scenarios in which apex predators are allowed to migrate into the environment when their densities fall below a threshold (<250 individuals). We therefore exclude the intrinsic population dynamics of specialist apex predators from further consideration.

When apex predators consume only mesopredators, the probability of coexistence increases by 3.4%, while the probability of extinction decreases by 4.1%. The dominant period of prey population oscillations increases by 4.26 weeks, and the dominant period of mesopredator oscillations increases by 3.57 weeks (Figure 5). The necessary conditions for coexistence remain largely unchanged relative to the baseline scenario (Appendix B: Table B1, Figure 3).

When apex predators target both mesopredators and prey, the probability of a prey only state increases by 10.5% in the presence of generalist apex predators with a corresponding reduction in the probability of the coexistence (−6.6%) and extinction (−3.9%) states. The dominant period of prey population oscillations is significantly higher (+34.86 weeks), while changes in mesopredator dynamics are comparable to those observed under specialist apex predator scenarios (+4.35 weeks). The necessary conditions for coexistence between mesopredators and prey shift towards higher overall mesopredator birth rates relative to specialist apex predator scenarios (i.e., scenarios where apex predators target only mesopredators).

### Varying the target of the superpredator

When superpredators consume mesopredators, the marginal probability of coexistence does not change relative to the baseline model, while the probability of a prey-only state increases slightly (+4.8%) and the probability of extinction decreases (–4.1%). When superpredators consume only prey, the probability of coexistence declines sharply (–30.7%) and the probability of extinction increases substantially (+30.9%; Figure 5; Appendix B: Table B1).

The inclusion of a generalist superpredator that consumes both mesopredators and prey leads to a small increase in the probability of the prey-only state (+4.9%), accompanied by modest reductions in the probabilities of coexistence (–3.1%) and extinction (–1.9%). Across all scenarios, the necessary conditions defining prey-only, coexistence, and extinction states remain consistent, regardless of which agents (or combinations of agents) are targeted by superpredators (Figure 4; Appendix B: Table B2).

**Figure 4:**
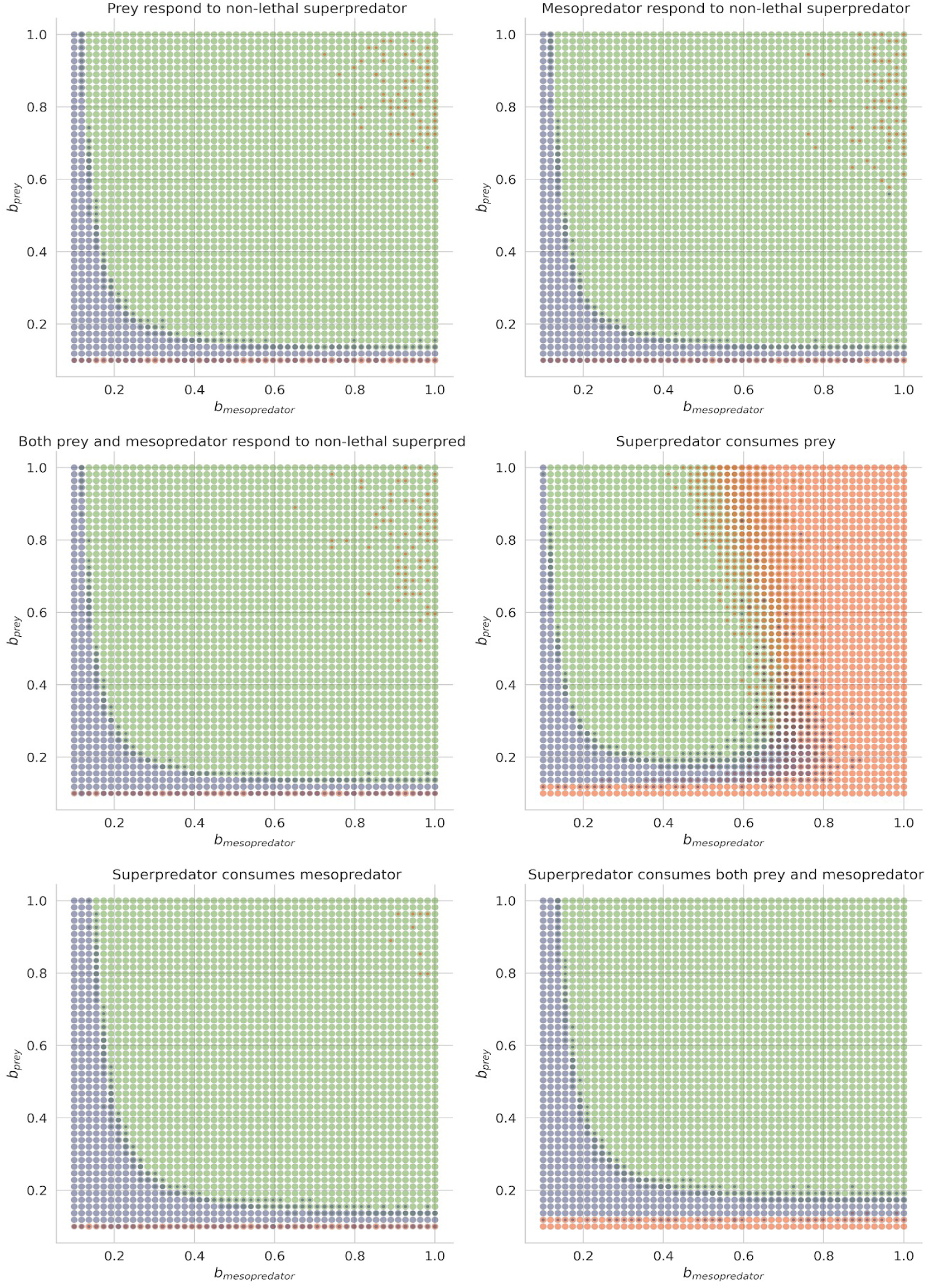
Outcomes of 25 repeated simulations across varying prey birth rates (bprey) and mesopredator birth rates (bmesopredator). (a) Scenarios in which superpredators replace apex predators by targeting prey, mesopredators, or both trophic levels. (b) Scenarios comparing lethal and non-lethal superpredators. Colours show the proportion of realisations resulting in Prey Only (blue), Coexistence (green), and Extinction (orange) across the parameter space.

The dominant period of both prey and mesopredator population oscillations decreases slightly when superpredators target prey. In contrast, when superpredators target mesopredators, the dominant period of oscillations for both groups increases slightly. The presence of generalist superpredators leads to a pronounced increase in the dominant period of prey population oscillations (+ 36 weeks) and a more modest increase in mesopredator oscillations (+7.08 weeks; Figure 5; Appendix C).

**Figure 5:**
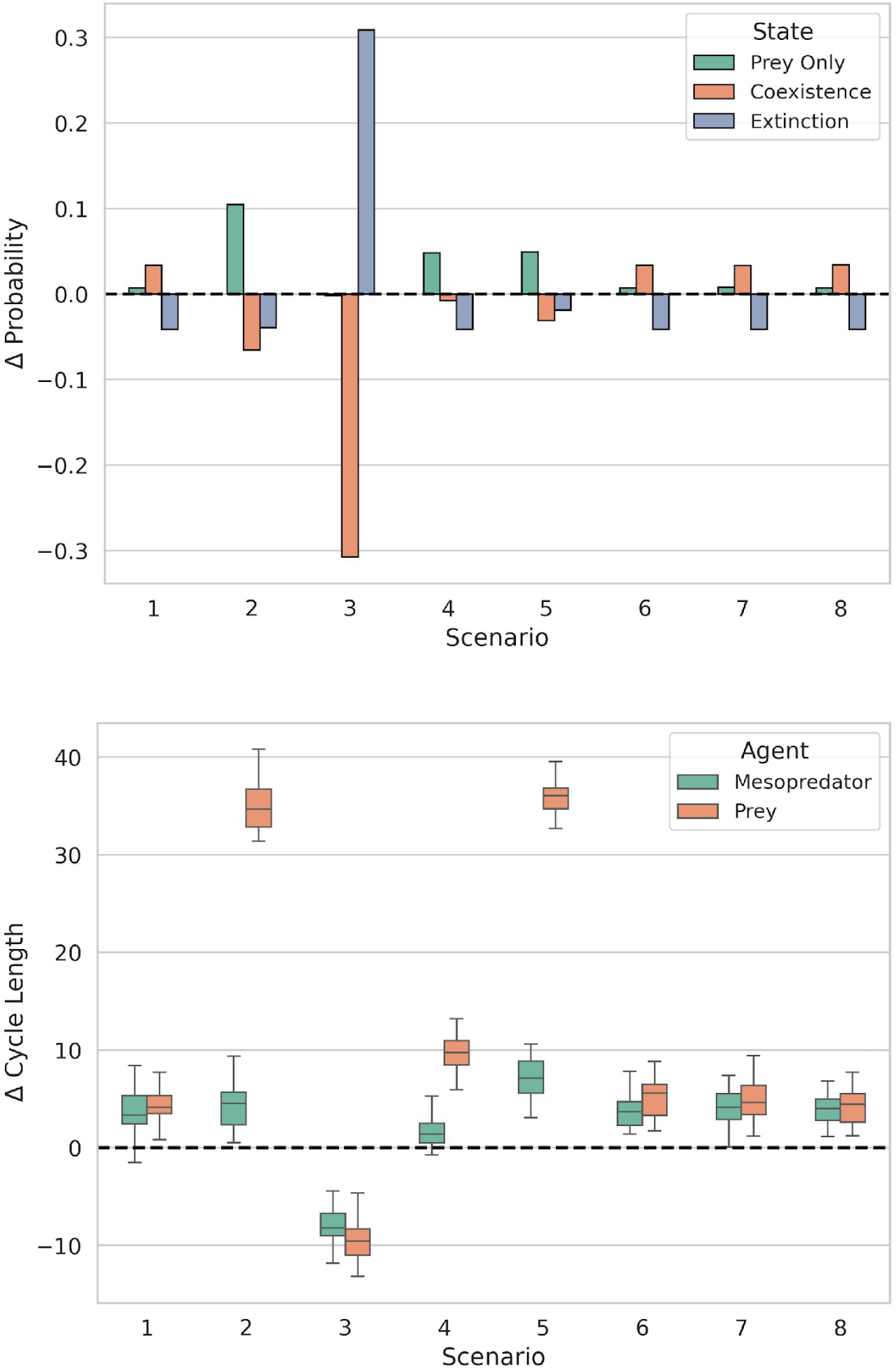
Change in probability of prey, only coexistence and extinction outcomes (top) and change in period of population oscillations (bottom) of each scenario defined in table 1 relative to the baseline scenario with only mesopredators and prey.

### Comparing lethal and non-lethal superpredators

Non-lethal superpredators produced similar effects on system dynamics regardless of whether mesopredators or prey responded to them behaviourally. The probability of coexistence increased slightly, accompanied by a corresponding decrease in the probability of extinction (Figure 5; Appendix B: Table B3). The necessary conditions defining coexistence, prey-only, and extinction states remained unchanged relative to the baseline scenario (Appendix B: Table B2). In addition, non-lethal superpredators exerted a small positive effect on the dominant period of both mesopredator and prey population oscillations (Appendix C).

## Discussion

This study aimed to explore how interactions with human superpredators alter predator-prey dynamics using an agent-based modelling approach. Unlike previous work, our model incorporates and compares both the consumptive (lethal) and non-consumptive (behavioural) effects of predators on prey (Table 1).Our results demonstrate that lethal superpredators alter the dynamics of mesopredator and prey populations to a greater extent than apex predators. The choice of trophic level targeted by the superpredator does not change model outcomes but does alter the population dynamics of prey agents (Figure 4 and 5). In contrast, non-lethal superpredators had minimal impact on the population dynamics of mesopredators and prey, even when they elicited anti-predator responses (Figure 5).

### Impact of replacing apex predators with superpredators

Humans have hunted predators to extinction in many ecosystems (e.g., Jackson et al. 2001). The removal of apex predators has often been intentional, motivated by perceived threats to livestock, competition for wild prey, or fears of danger. Over time, the systematic decline of apex predators has led to a progressive expansion of human influence downward through food webs, as people began targeting mesopredators and herbivores. Humans have effectively replaced apex predators as top-down regulators of mesopredator and prey populations through hunting, fishing, and trapping (Pauly et al. 1998). However, the extent and mechanism of humans replacing apex predators remains poorly understood.

In our model, specialist apex predators have a much smaller effect on mesopredator and prey population dynamics than specialist superpredators. This likely reflects the low encounter success and low population densities of apex predators. Because apex predators roam across vast areas, their local influence can be limited. In contrast, human hunters and fishers can harvest large numbers of animals from small areas with high efficiency, leading to much stronger local impacts (Estes et al. 2011). Generalist apex predators had stronger effects on model outcomes and population dynamics than specialist apex predators, but still substantially weaker effects than lethal superpredators (Figure 5).

Trophic downgrading and the replacement of apex predators with superpredators reshape energy flow, species interactions, and ecosystem stability (Estes et al. 2011). Mesopredator release following the loss of apex predators has been known to cause population collapses of prey species in some ecosystems (Ritchie & Johnson, 2009). For example, the extirpation of large shark species on the east coast of North America had a cascading positive effect on the population of Cow Nose rays (*Rhinoptera bonasus*), the increased predation pressure on lower trophic levels lead to the eventual collapse of scallop fisheries in the region (Myers et al., 2007). In the current model, when specialist apex predators were present, mesopredator populations were only weakly suppressed (coexistence probability increased by just 3.4%), whereas lethal superpredators produced far larger shifts in community state (up to −30.7% coexistence when targeting prey directly), consistent with the stronger demographic control humans exert in real ecosystems.

On the other hand, mesopredator suppression by superpredators may promote coexistence between mesopredators and prey. Similar patterns occur in nature. For example, heavy hunting pressure on wolves has allowed human harvest to regulate mesopredator populations more strongly than wolves alone ever did (Ripple et al., 2014). Intense human harvesting has also suppressed mid-trophic predators in marine systems, such as large demersal fishes on the Scotian Shelf and throughout the Mediterranean Sea (Coll et al., 2012; Frank et al., 2005). These ecological changes have substantial social and economic consequences. The removal or replacement of apex predators by human superpredators disrupts services such as pest control, carbon storage, and fisheries productivity, which can impose long-term economic costs (Burgos et al., 2024).

### Effects of superpredator target selection

Human superpredators vary in which species they target across both time and space (Treves & Naughton-Treves 1999), and the species they do target are harvested with remarkable efficiency (Darimont et al., 2015). In our model, the choice of target for lethal superpredators significantly influenced system dynamics. When superpredators consume mesopredators, exclusively or in addition to prey, the probability of mesopredator – prey coexistence increased. Correspondingly, the probability of extinction for all agents decreased while the probability of the prey only state remained largely unchanged. This pattern indicates a stabilising effect of superpredators on predator–prey dynamics. Similar stabilisation occurs in nature when strong human pressure suppresses dominant mid-level predators and reduces competitive or consumptive release. For example, targeted culling of coyotes can temporarily reduce their impact on smaller carnivores and prey, which can increase local biodiversity in the short term (Prugh et al., 2009). Likewise, reductions in mesopredatory fish through intense fishing have allowed prey fish to persist in some marine systems (Heithaus et al., 2008). However, coexistence under mesopredator-focused superpredators required higher birth rates for both mesopredators and prey. This suggests that both groups face stronger demographic challenges under intense human predation. Elevated reproductive rates compensate for increased mortality, a pattern also seen in heavily hunted ungulates and marine fish stocks (Jørgensen et al., 2007).

When superpredators consume only prey, the probability of coexistence between mesopredators and prey decreased significantly while the probability of extinction increased significantly. In contrast, specialist superpredators targeting exclusively mesopredator had limited effect on model outcomes. . Prey populations also showed fewer oscillations under direct superpredator pressure (Figure 5). This damping likely resulted from strong top-down forcing that prevented the prey populations from increasing rapidly. Heavy and consistent predation can truncate population cycles by removing individuals before density-dependent processes unfold. Similar effects occur when human hunting suppresses cyclic dynamics in species such as snowshoe hares, moose, and cod (Frank et al., 2007; Krebs et al., 2001). These patterns mirror empirical cases where high human offtake from shared prey pools leads to prey declines and secondary collapses of predators that depend on them (Ripple et al. 2014). Such dynamics highlight the far-reaching consequences of human target choice and harvesting intensity for the stability of ecological communities.

### Effect of lethal vs. non-lethal superpredators

In addition to hunting, fishing and other lethal interactions, humans often interact with wild animals in non-lethal ways, such as through tourism or recreation (Boyle & Samson, 1985). The risk-disturbance hypothesis posits that animals should respond to benign humans in the same way they do to any other predator (Frid & Dill, 2002). In our model, non-lethal superpredators elicited prey behavioural responses from their “target” agents without consuming them. These non – lethal interactions had a minimal impact on the system’s state compared to their lethal counterparts. Our findings suggest that even when mesopredators and prey respond to non-lethal superpredators, these interactions may not significantly affect their populations.

Our results suggest that behavioural responses of targetted species alone may not destabilise populations unless they impose significant energetic or opportunity costs. Reduced foraging, increased movement, and elevated stress in response to higher predation pressure may lower survival or reproduction in real systems (Clinchy et al. 2004; Zanette et al. 2011). Such non-consumptive effects have been shown to reduce prey fitness in many taxa, including ungulates exposed to trekkers (Neumann et al., 2009) and marine mammals disturbed by vessel traffic (Williams et al., 2006). If these costs were larger in our model, non-lethal superpredators would likely have stronger effects. However, evidence for these potential fitness costs translating into demographic effects is limited and disputed (Sheriff et al., 2020). All else being equal, lethal superpredators exert a much greater impact on predator - prey population dynamics (see Dsouza et al., 2025).

### Limitations and future directions

In our simulations, apex predators were unable to persist in our simulations regardless of birth rate, lethality and handling limit values chosen. This result contrasts with many mean – field models of tritrophic interactions which do show coexistence regimes both when behavioural anti-predator responses are incorporated (Hastings & Powell, 1991; Upadhyay & Rai, 1997), and when they are not (Verma et al., 2021). In nature, apex predators also face strong constraints. Apex predators require much larger home ranges than lower trophic levels because of higher energy demand and hunting low success. They must search widely to encounter enough prey to meet their metabolic needs (Carbone et al., 2007; Carbone & Gittleman, 2002).

The lack of an apex predator – mesopredator – prey coexistence regime in our model indicates that top-down control alone does not allow for a stable trophic network. Instead, stable coexistence requires a complex balance of top down and bottom-up regulation (Rooney et al., 2006). Our results could also be attributed to our choice of discrete time simulation which can promote instability and extinction even when continuous time models are stable (Hastings, 1990). A larger spatial grid (>100 cells per side) might allow for coexistence by providing the required space for an apex predator to exist. An additional trophic level could also provide stability by adding bottom-up feedback. However, these additions are computationally costly. To facilitate our comparison of apex predators and superpredators, we instead implemented apex predator migration which ensures that apex predators were present throughout our simulations.

While our model provides valuable insights, it assumes homogeneous space without environmental variability or habitat complexity, which has a large impact on how wild animals respond to predators (Wirsing et al., 2021). Simplified agent behaviours may not capture the full range of real-world interactions, and although we account for conspecific competition through carrying capacity, we do not explicitly model resource distribution or abundance. A further limitation is that our model includes only a single species per trophic level, whereas real food webs involve multiple species with varying degrees of connectivity and dependency within and between trophic levels. The consumptive and non-consumptive effects of predators and humans may be substantially stronger or weaker in more complex food webs, where indirect effects, competitive release, and alternative prey can buffer or amplify trophic cascades (Polis et al., 2000; Terborgh et al., 2001).

Future research should integrate habitat heterogeneity, resource distribution, and environmental stochasticity into models to better reflect real-world ecosystems. Exploring the effects of spatial refuges and movement barriers could provide a deeper understanding of population dynamics (Mitchell & Lima, 2002). Expanding the range of behavioural responses, such as learning and memory, would enhance the model’s realism. Empirical studies are needed to test predictions in natural ecosystems affected by human activities, particularly to observe behavioural adaptations of prey and predators in response to superpredator pressures. Investigating socio-economic factors and incorporating human decision-making processes into ecological models could also improve their accuracy and applicability (Darimont et al. 2023).

## Conclusions

We demonstrate that lethal human “superpredators” play a pivotal role in shaping predator-prey dynamics, influencing the likelihood of coexistence, extinction, and population stability. Both target selection and lethality of superpredators are critical factors determining ecosystem outcomes. Our model provides a useful framework for managers seeking to regulate hunting pressure and wildlife populations, for example by informing decisions about the number of permits to issue annually. While the model provides a valuable framework for understanding these interactions, future work should incorporate greater ecological complexity and validate findings with empirical data. By doing so, we can better predict and mitigate the impacts of human activities on natural ecosystems.

## Supporting information

Apexpredator model

Base model

Superpredator Model

## Acknowledgments

The author thanks Kartik Shanker, Maria Thaker, and Vishwesha Guttal for their valuable feedback on the manuscript and for helpful discussions on the underlying concepts. I would like to thank KS for access to computational resources. I would like to thank the Prime Minister’s Research Fellowship for funds to conduct this work.

## Author Contribution Statement

Shawn Dsouza: Conceptualization; Methods development/experimental design; Software development; Coding simulation; Model analysis; Data validation; Data visualization; Writing – original draft; Writing – review & editing.

