## Supplementary Materials for "Multi-trophic risk from human superpredators may alter predator-prey coexistence and population dynamics"

**Appendix A: Model parametrisation**

We tested various parameter combinations to identify a parameter space that yielded ecologically interesting dynamics for hypothesis testing. We evaluated 50 evenly spaced values for each parameter independently while keeping the remaining parameters fixed (see Table 2 for default parameter values). For models where apex predators consume prey, we did not implement the migration rule during testing.

*Effect of maximum energy (handling limit) Emax on specialist apex predator dynamics*

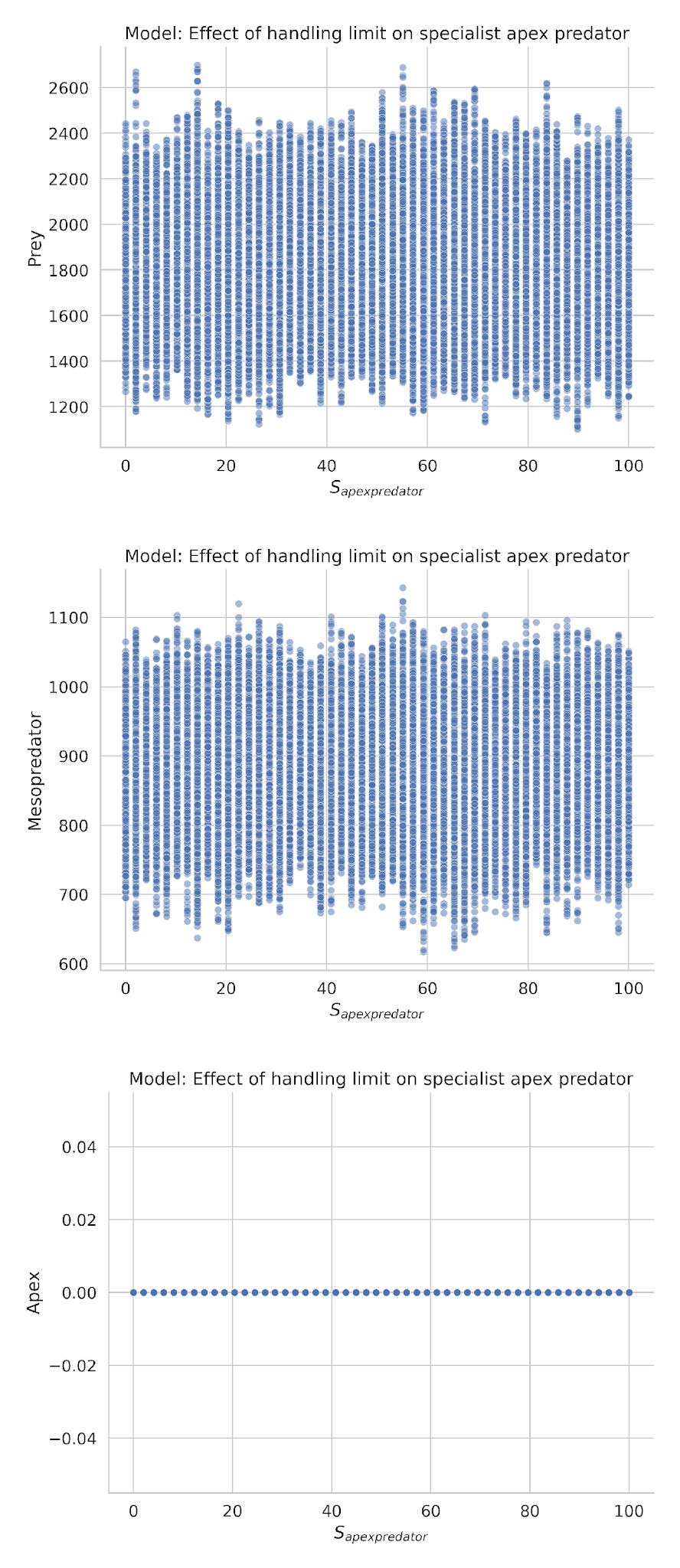

**Figure A1:** Equilibrium (T = [400, 1000]) populations of mesopredators and prey for 10 realizations across varying levels of apex predator handling limit. We varied apex predator lethality between 0 and 100. We found no parameters where specialist apex predators persisted to the end of the realization.

*Effect of varying apex predator (specialist) birth rate on agent dynamics*

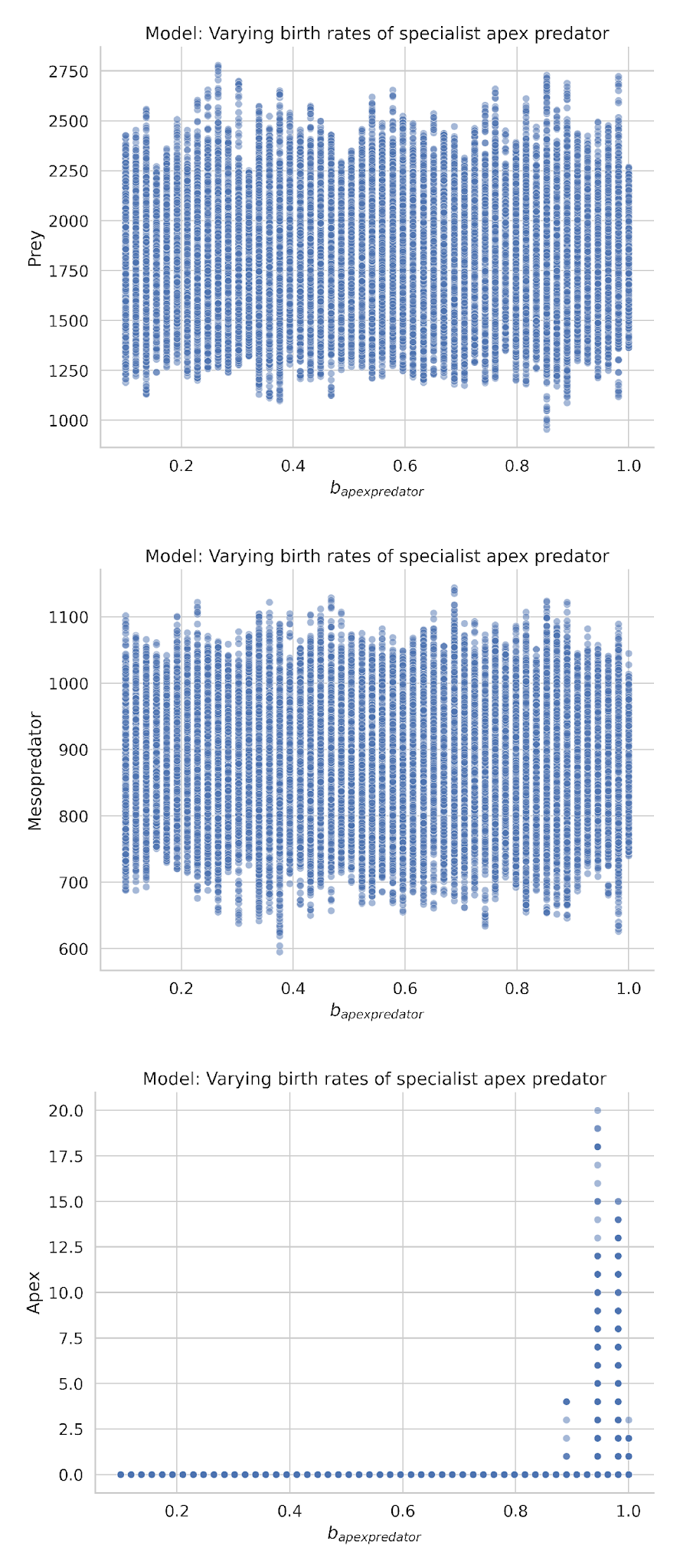

**Figure A2:** Equilibrium (T = [400, 1000]) populations of apex predator, mesopredators and prey for 10 realizations across varying levels of apex predator birth rates. We varied apex predator birth rates between 0 and 1.

*Effect of varying specialist apex predator lethality on agent dynamics*

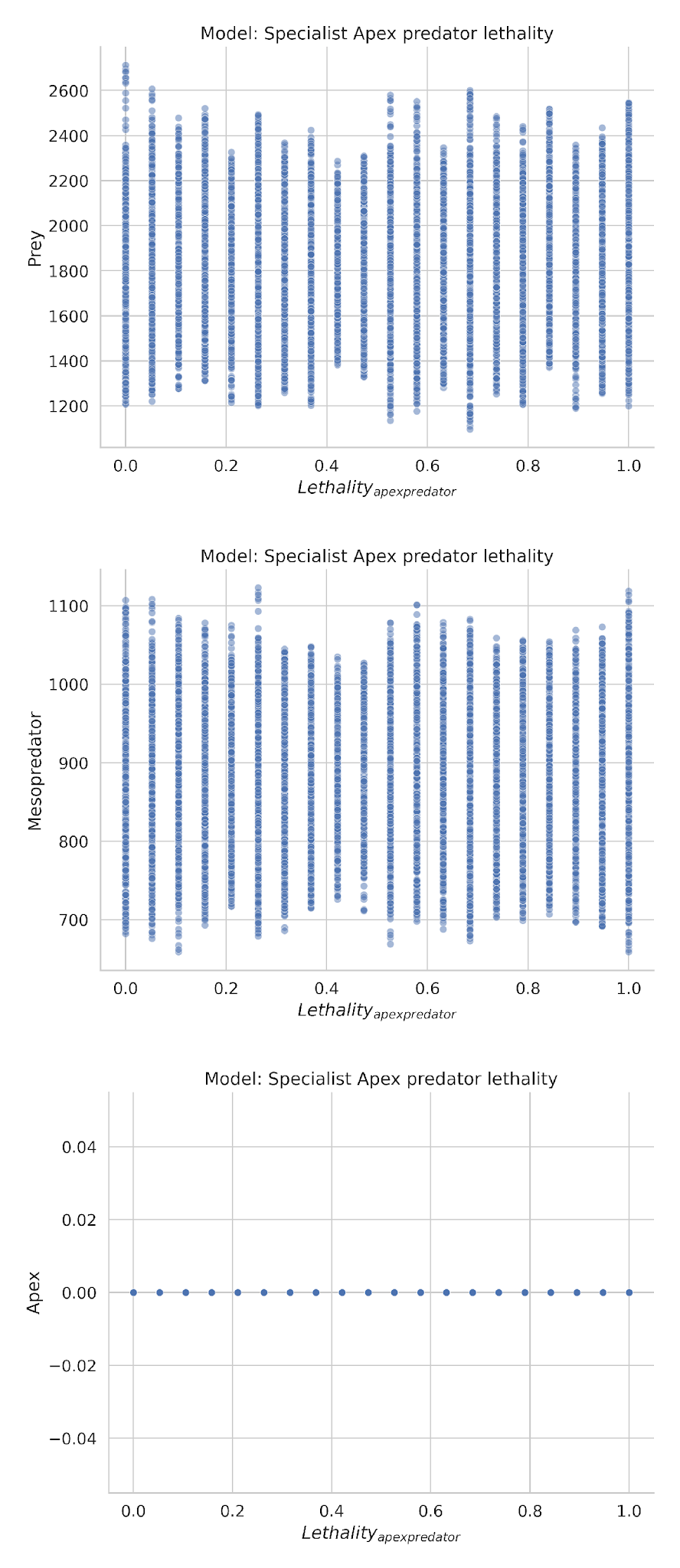

**Figure A3:** Equilibrium (T = [400, 1000]) populations of apex predator, mesopredators and prey for 10 realizations across varying levels of apex predator lethality. We varied apex predator lethality between 0 and 1. Once again, we found no parameters where apex predators persisted to the end of the realization.

*Effect of maximum energy (handling limit) Emax on generalist apex predator dynamics*

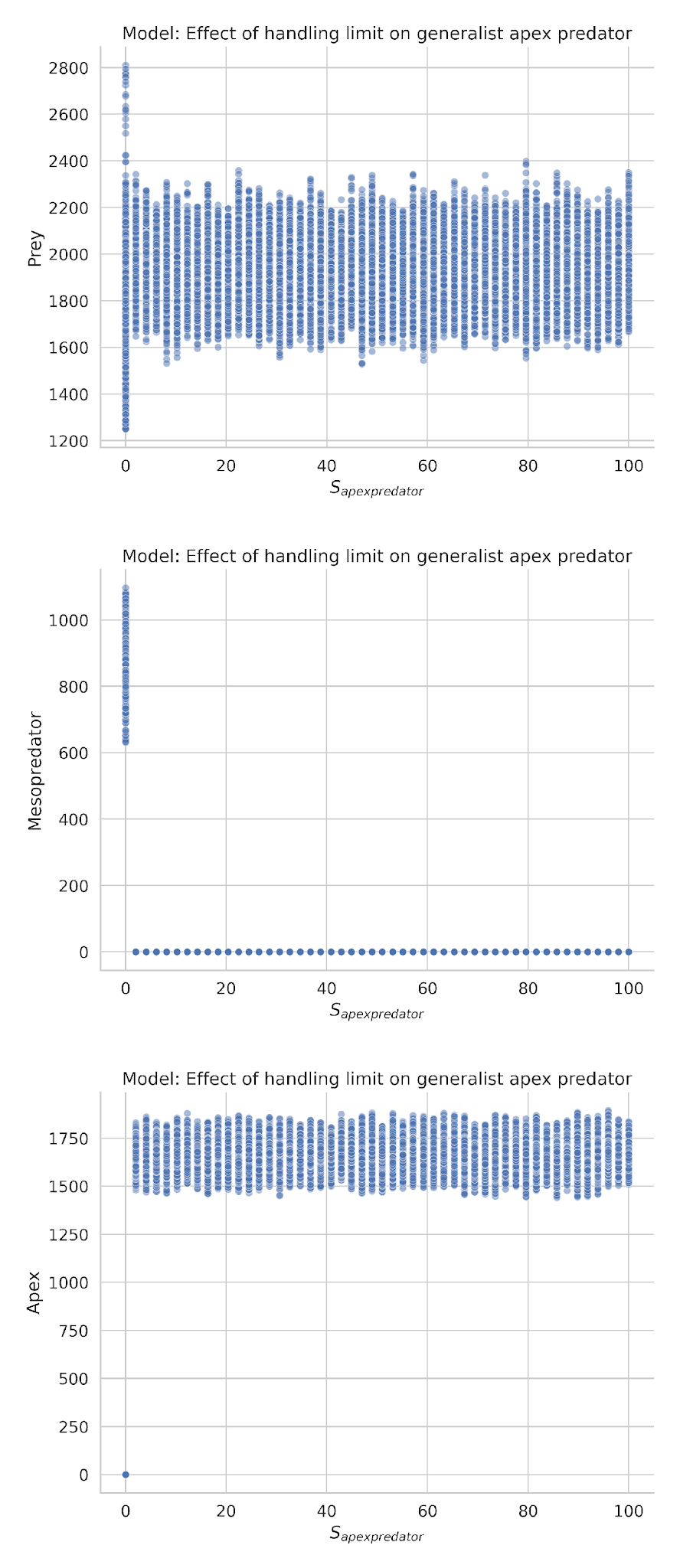

**Figure A4:** Equilibrium (T = [400, 1000]) populations of mesopredators and prey for 10 realizations across varying levels of apex predator handling limit. We varied apex predator lethality between 0 and 100.

*Effect of varying apex predator (generalist) birth rate on agent dynamics*

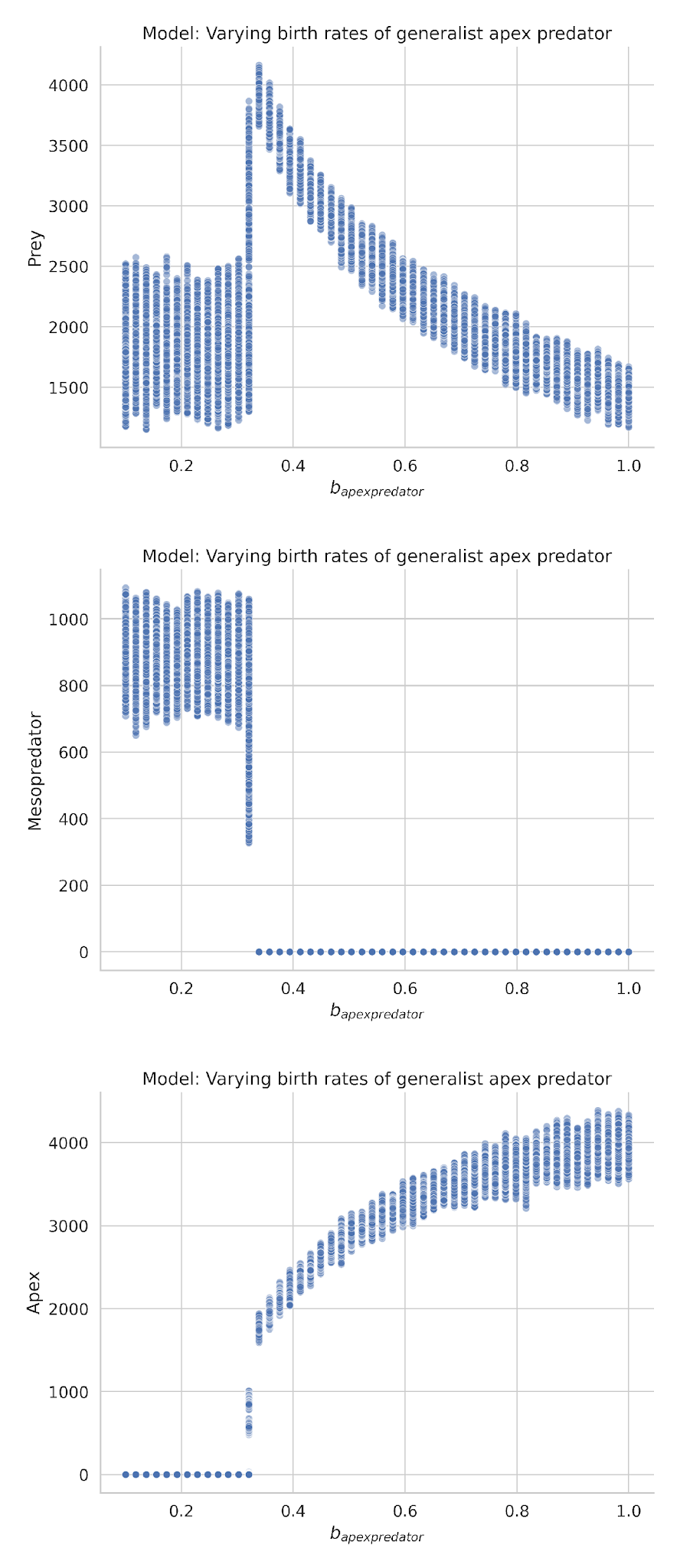

**Figure A4:** Equilibrium (T = [400, 1000]) populations of mesopredators and prey for 10 realizations across varying levels of apex predator birth rate. We varied apex predator birth rate between 0 and 1.

*Effect of varying generalist apex predator lethality on agent dynamics*

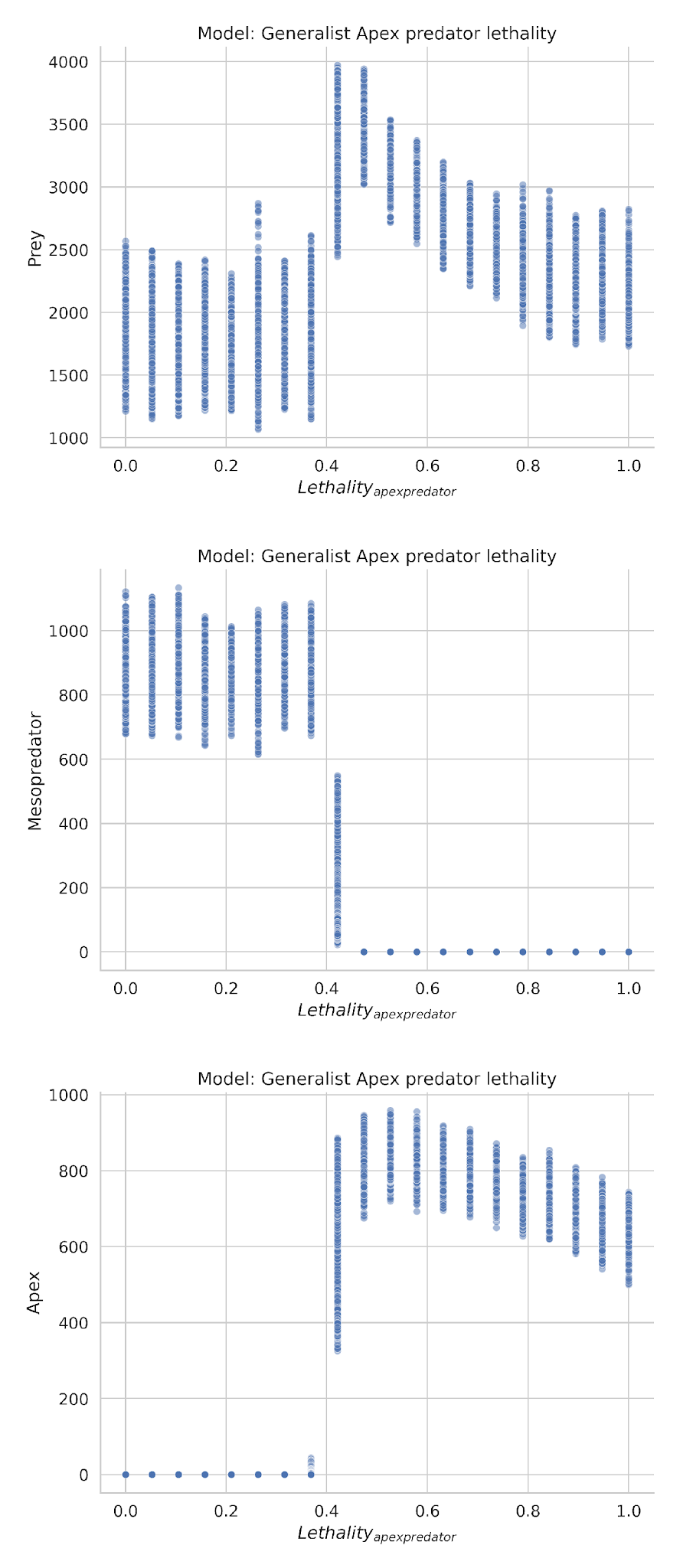

**Figure A5:** Equilibrium (T = [400, 1000]) populations of mesopredators and prey for 10 realizations across varying levels of apex predator lethality. We varied apex predator lethality between 0 and 1.

*Effect of maximum energy (handling limit) Emax* *on mesopredator dynamics*

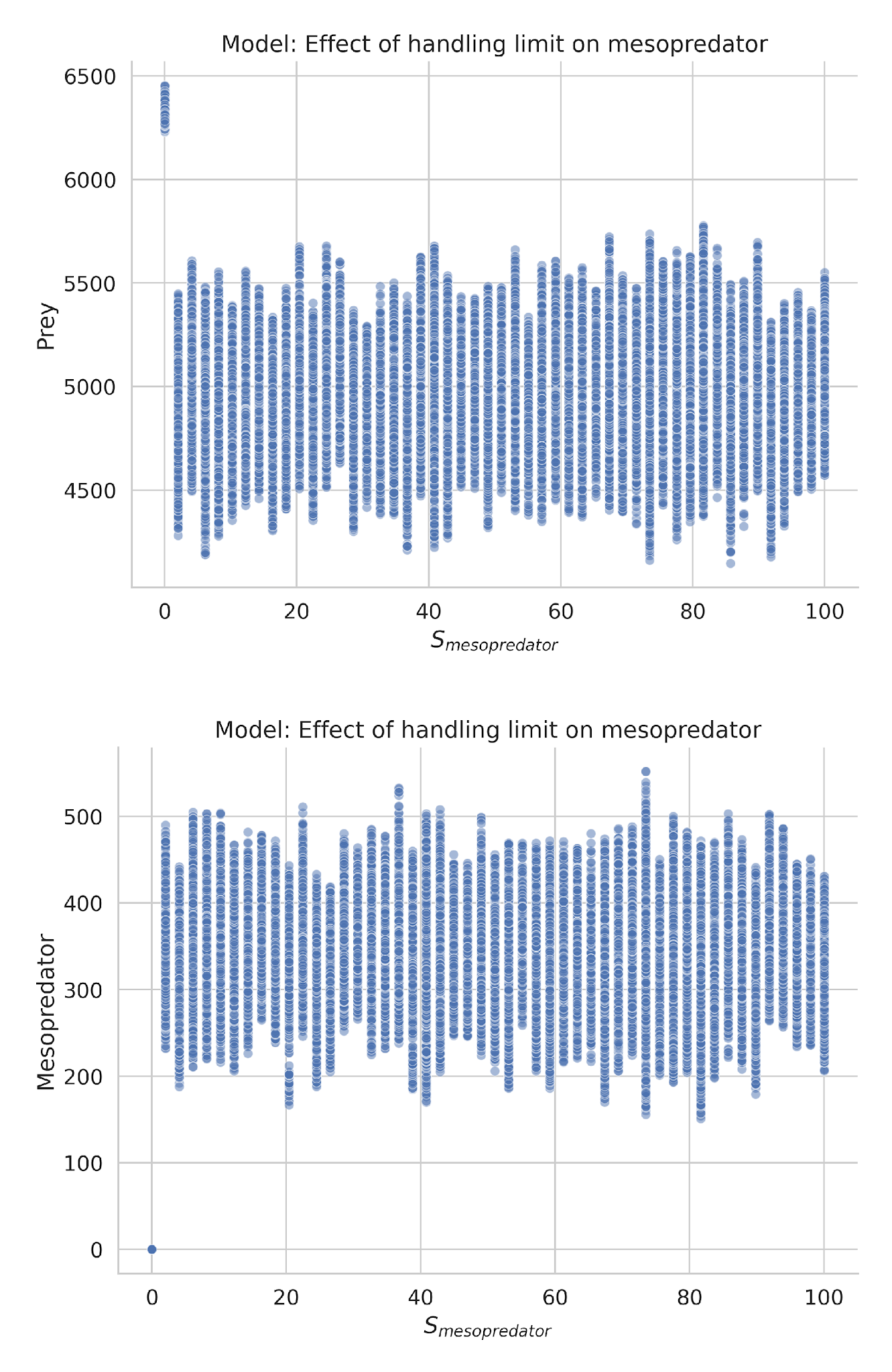

**Figure A6:** Populations of predator and prey at equilibrium (T = [400,1000]) for 10 realizations with varying maximum energy (Emax) of mesopredator. We varied the maximum energy that mesopredators can gain between 1 and 100 and no discernible change in model dynamics.

*Effect of varying mesopredator lethality on agent dynamics*

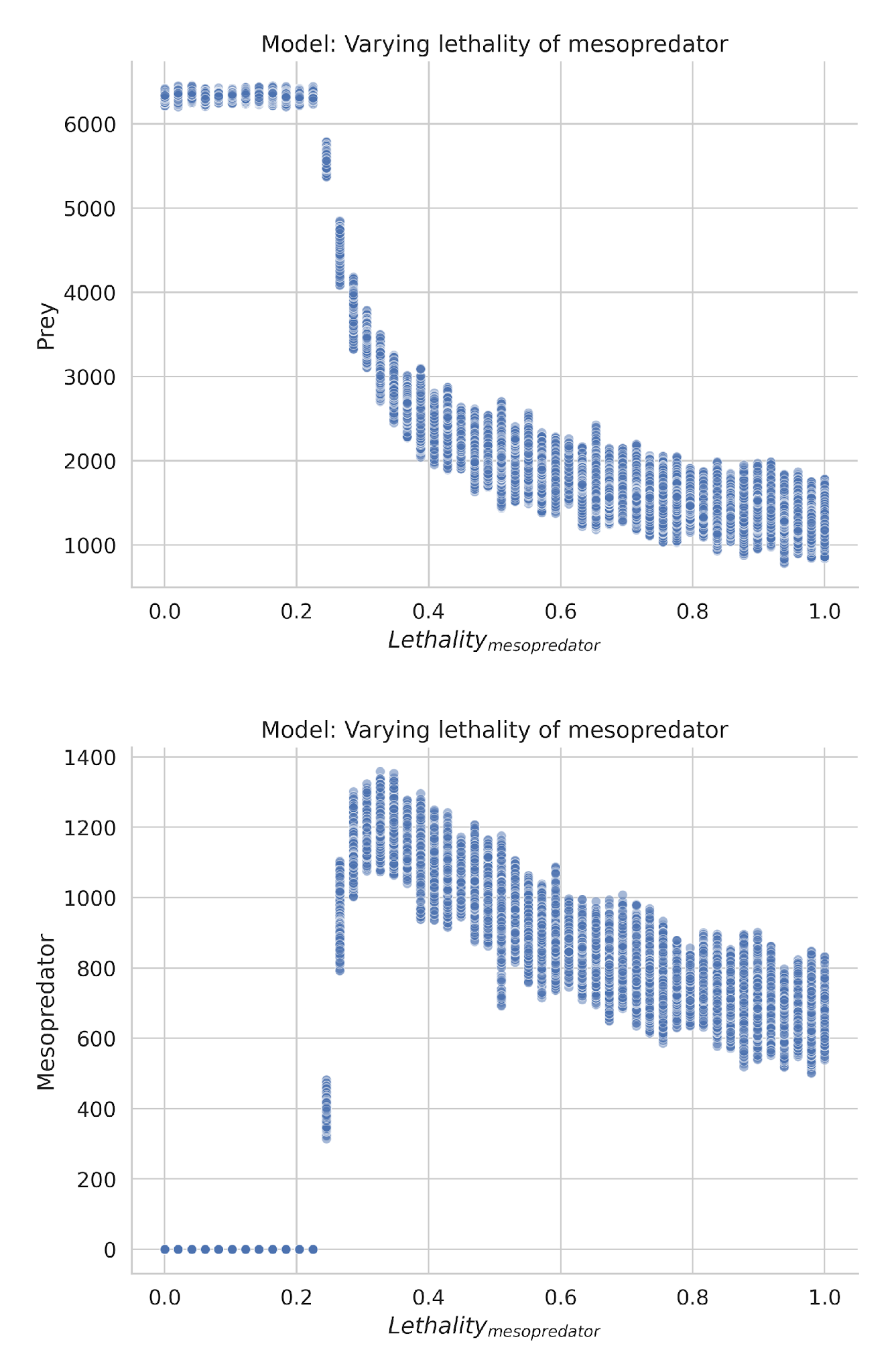

**Figure A7:** Equilibrium (T = [400, 1000]) populations of mesopredators and prey for 10 realizations across varying levels of mesopredator lethality. We varied apex predator lethality between 0 and 1. We observe a transcritical bifurcation around lmesopredator=0.26, which shift the system from a prey only state to a coexistence state. Beyond lmesopredator=0.3 the system appears stable.

*Effect of lattice size (L2) and local saturation of prey (S) on equilibrium prey populations*

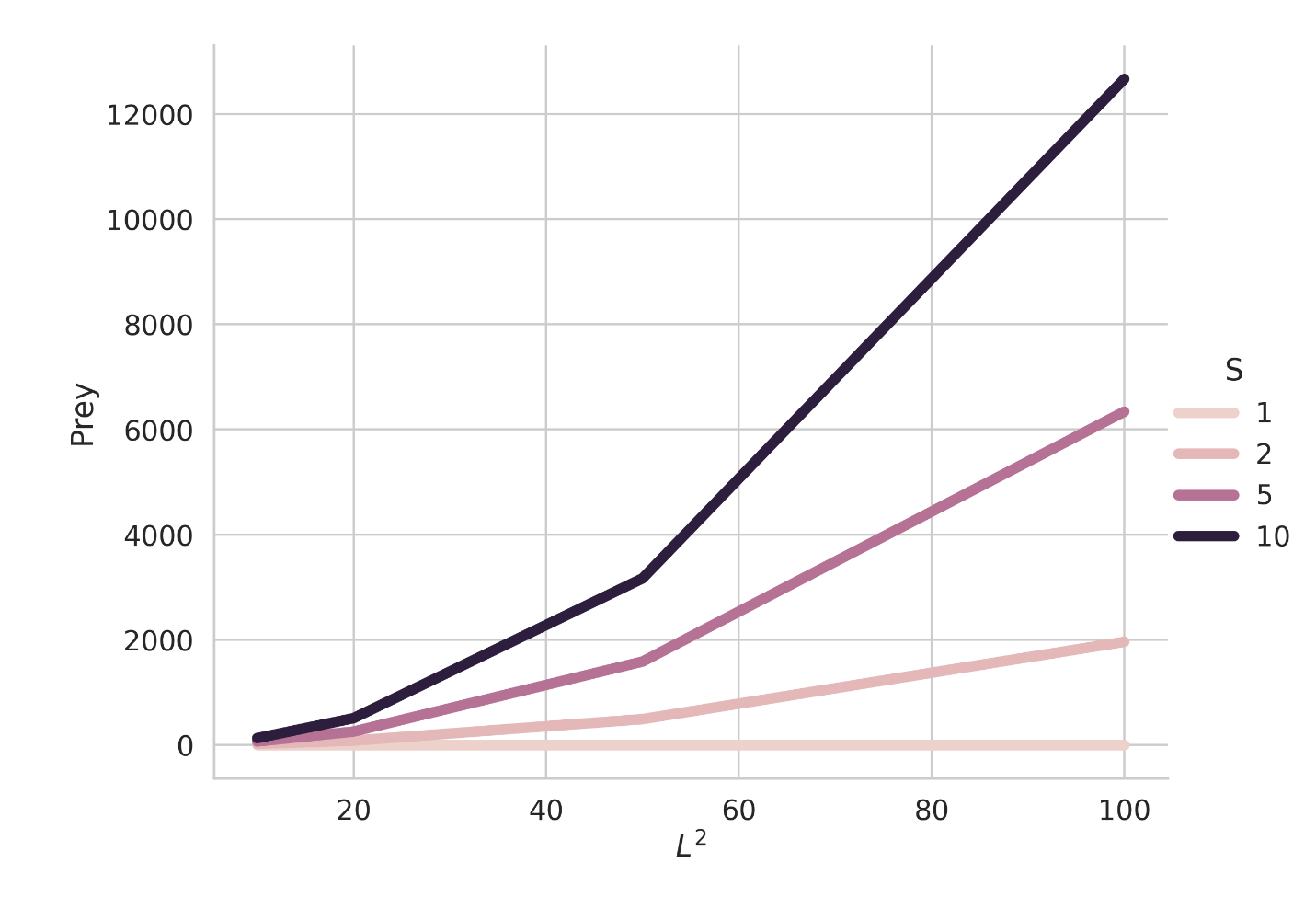

**Figure A8:** Equilibrium (T = [400, 1000]) prey populations averaged over 10 realizations with varying lattice size (*L2*) and local prey saturation (*S*). The equilibrium population of prey increases linearly with lattice size and local saturation. Based on these results, we selected *L2*  =100 and *S*=5, where the average equilibrium population N400,1000¯=6300, as computational challenges arise beyond these values. This choice ensures a balance between ecological relevance and computational feasibility.

*Effect of starting density of prey, mesopredator superpredators and apex predators on agent dynamics.*

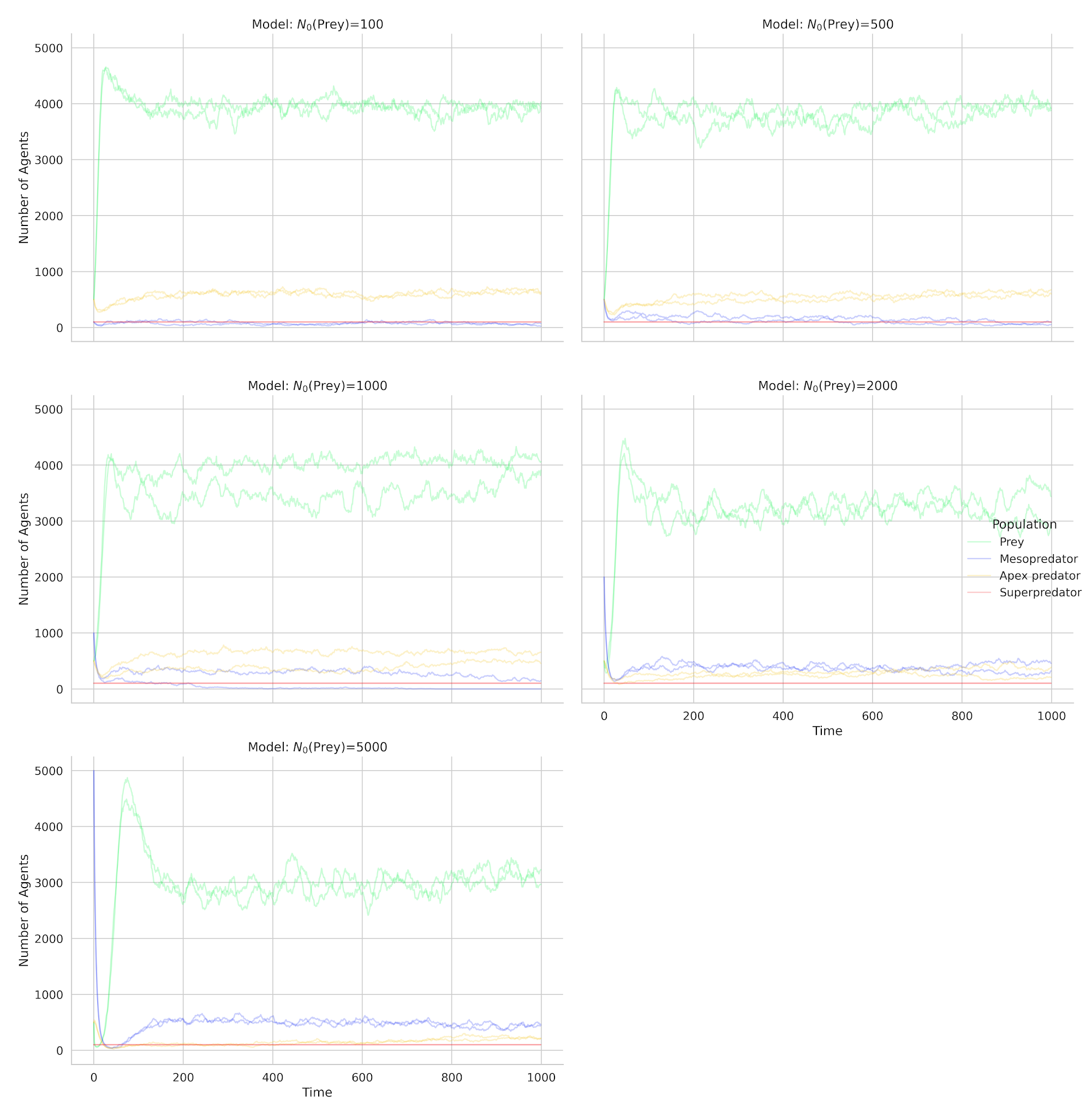

**Figure A9:** Population timeseries of 5 representative realizations at varying starting densities of prey. Prey starting densities had no effect on agent dynamics. We chose N0=500 for prey as that is closer to equilibrium.

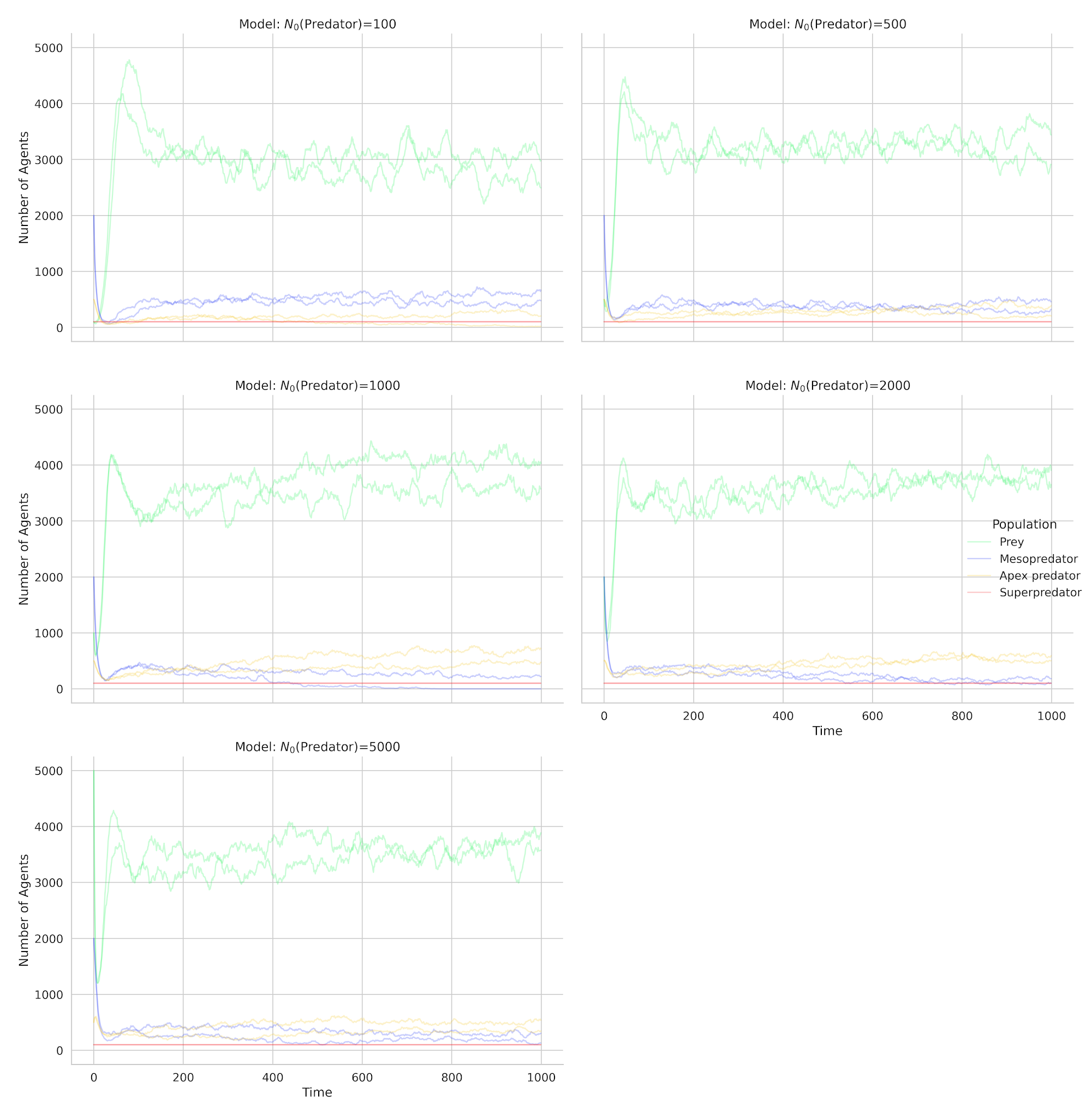

**Figure A10:** Population timeseries of 5 representative realizations at varying starting densities of mesopredator. Mesopredator starting densities had no effect on agent dynamics. We chose N0=500 for mesopredator as that is closer to equilibrium.

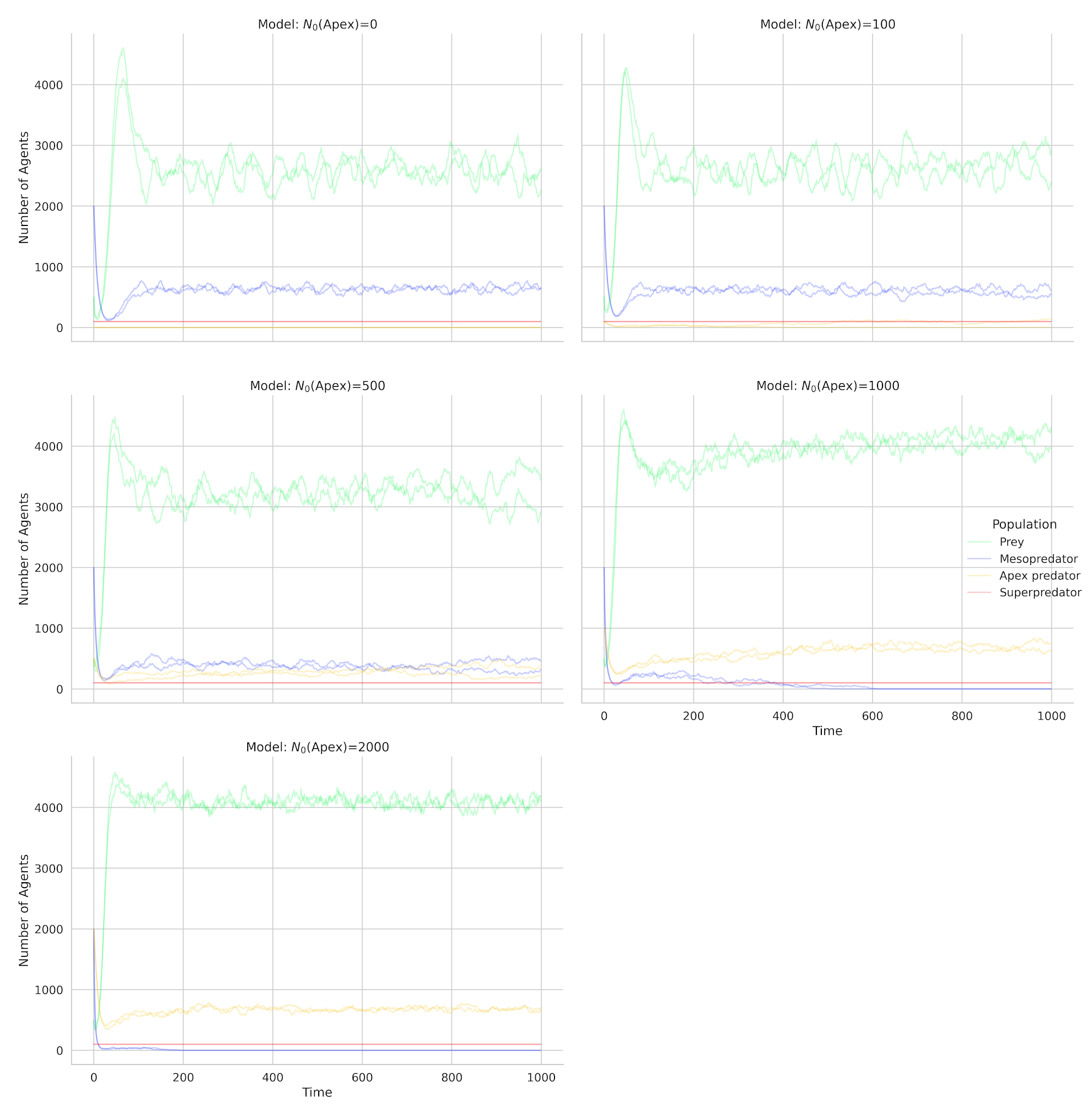
**Figure A11:** Population timeseries of 5 representative realizations at varying starting densities of apex predator. Apex predator starting densities had no effect on agent dynamics. We chose N0=500 for apex predator.

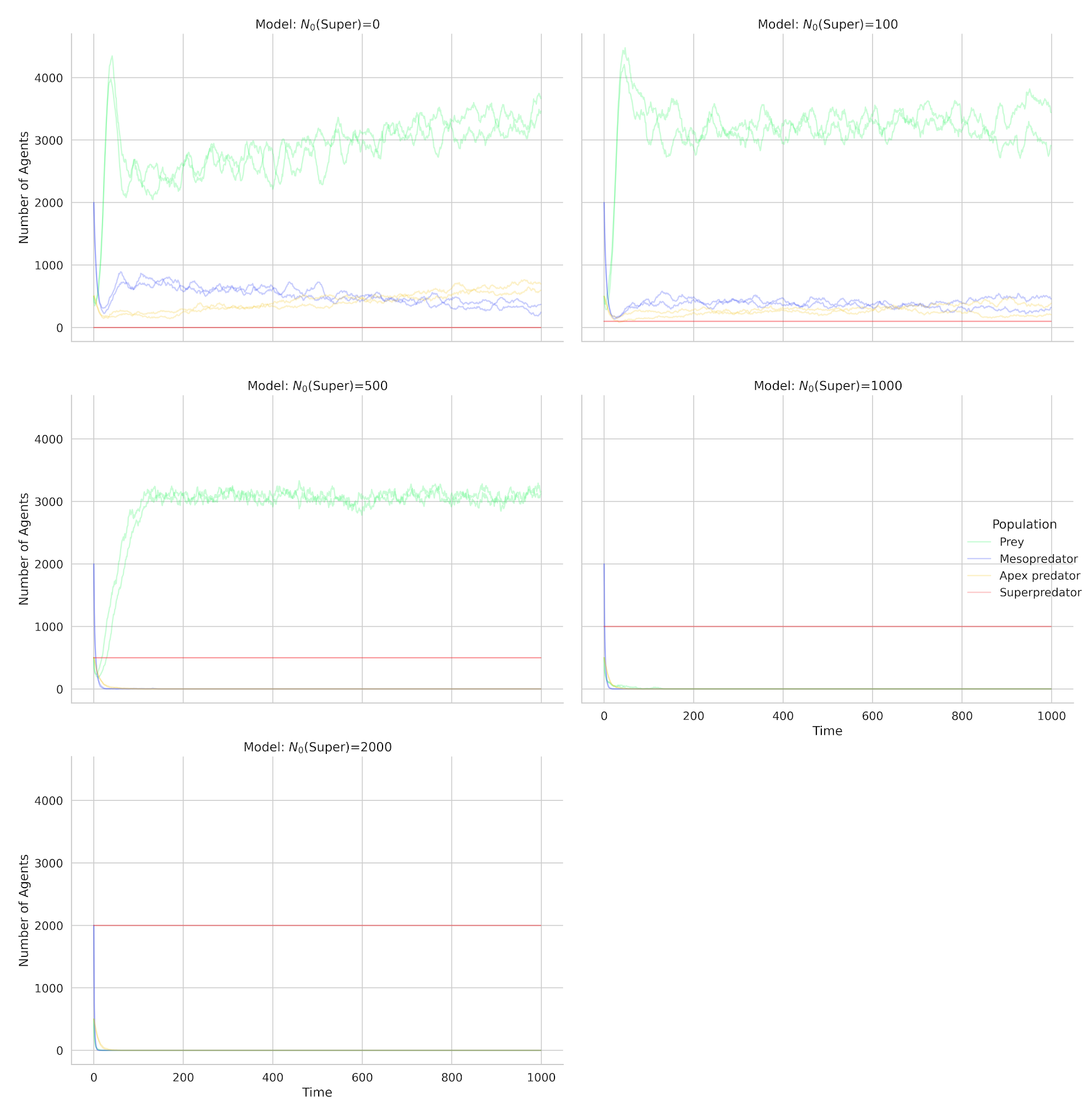
**Figure A12:** Population timeseries of 5 representative realizations at varying starting densities of superpredators We chose N0=100 for superpredators as that allows for coexistence of mesopredators and prey.

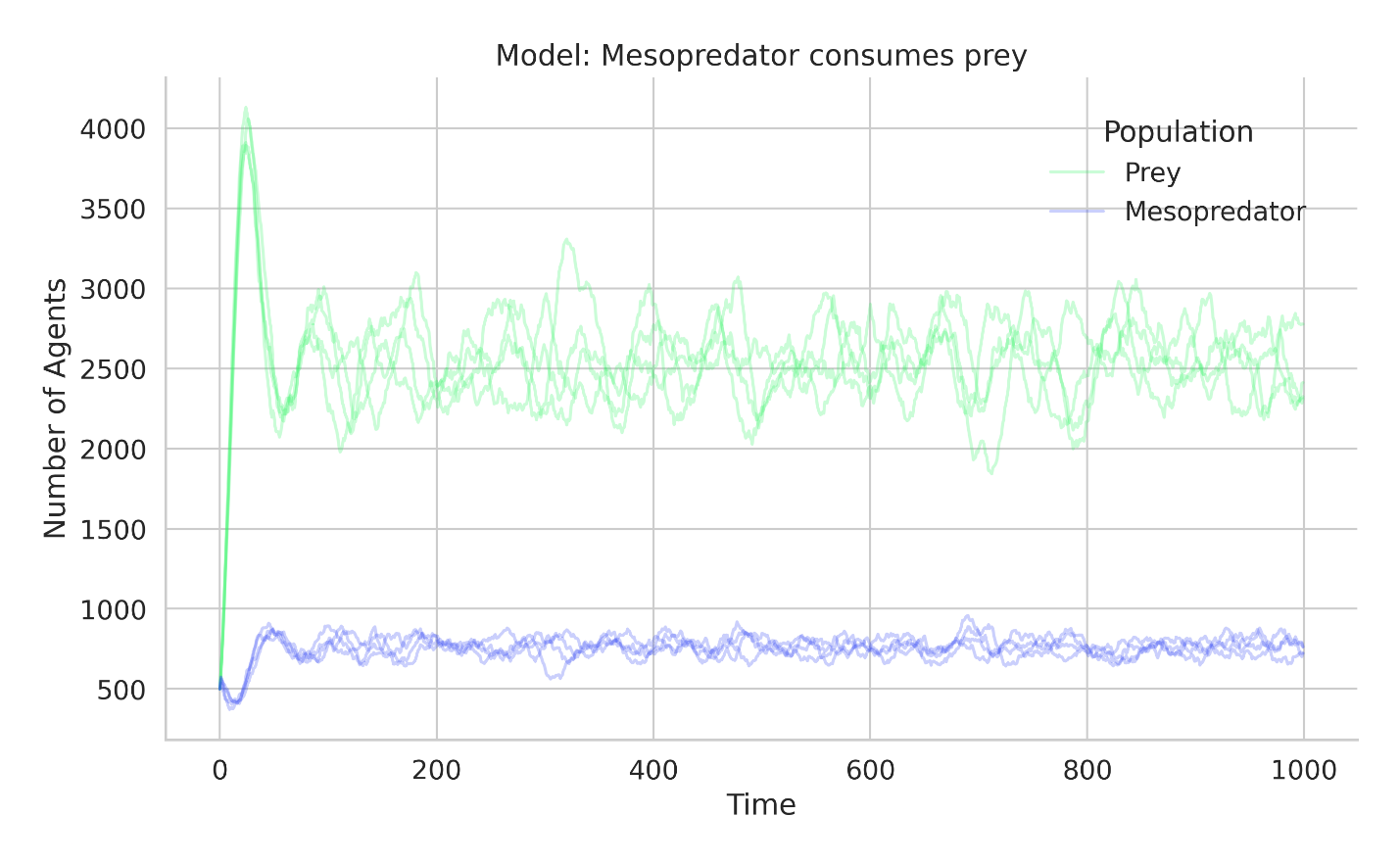

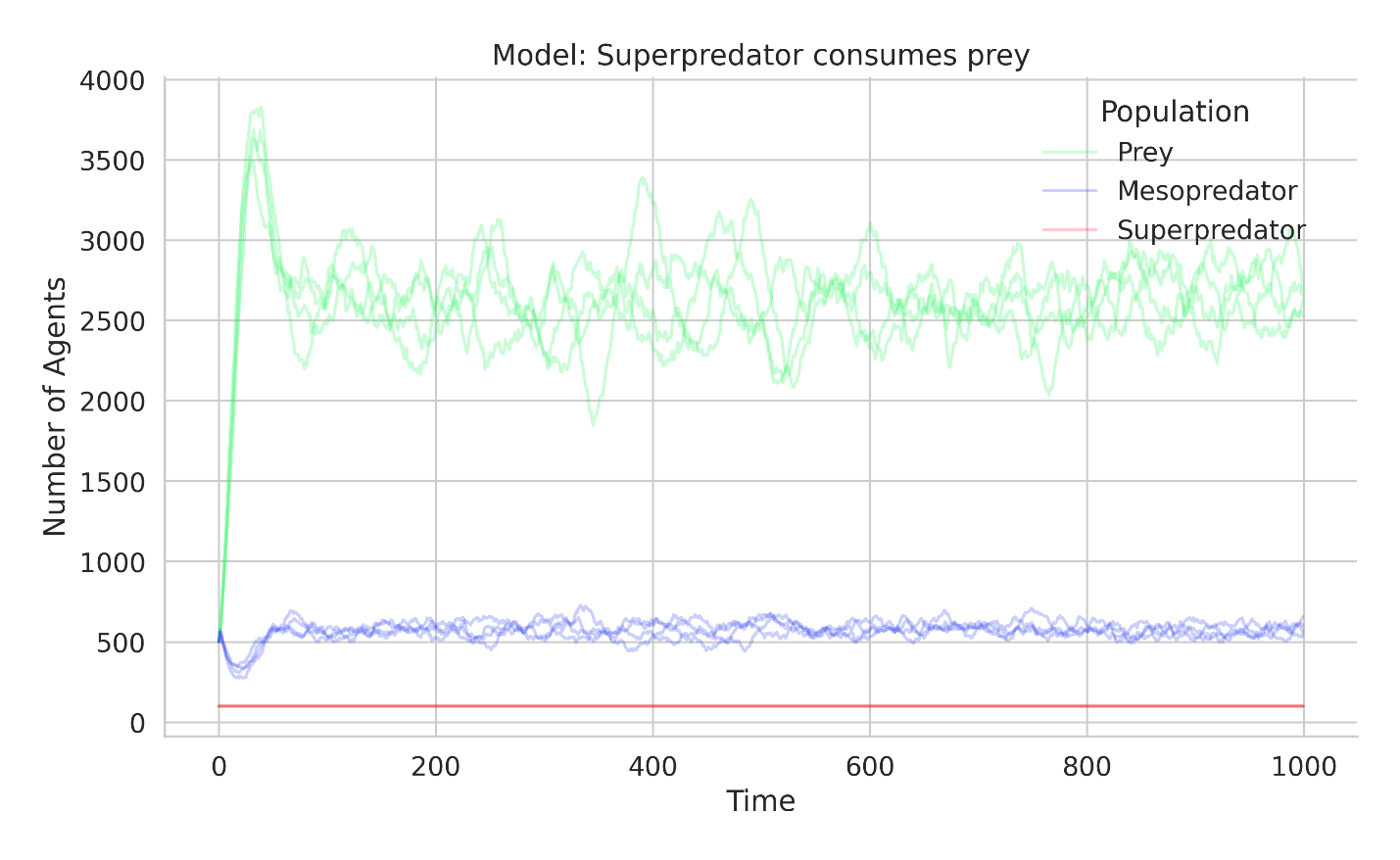

**Figure A13:** Selected examples of steady-state population timeseries.

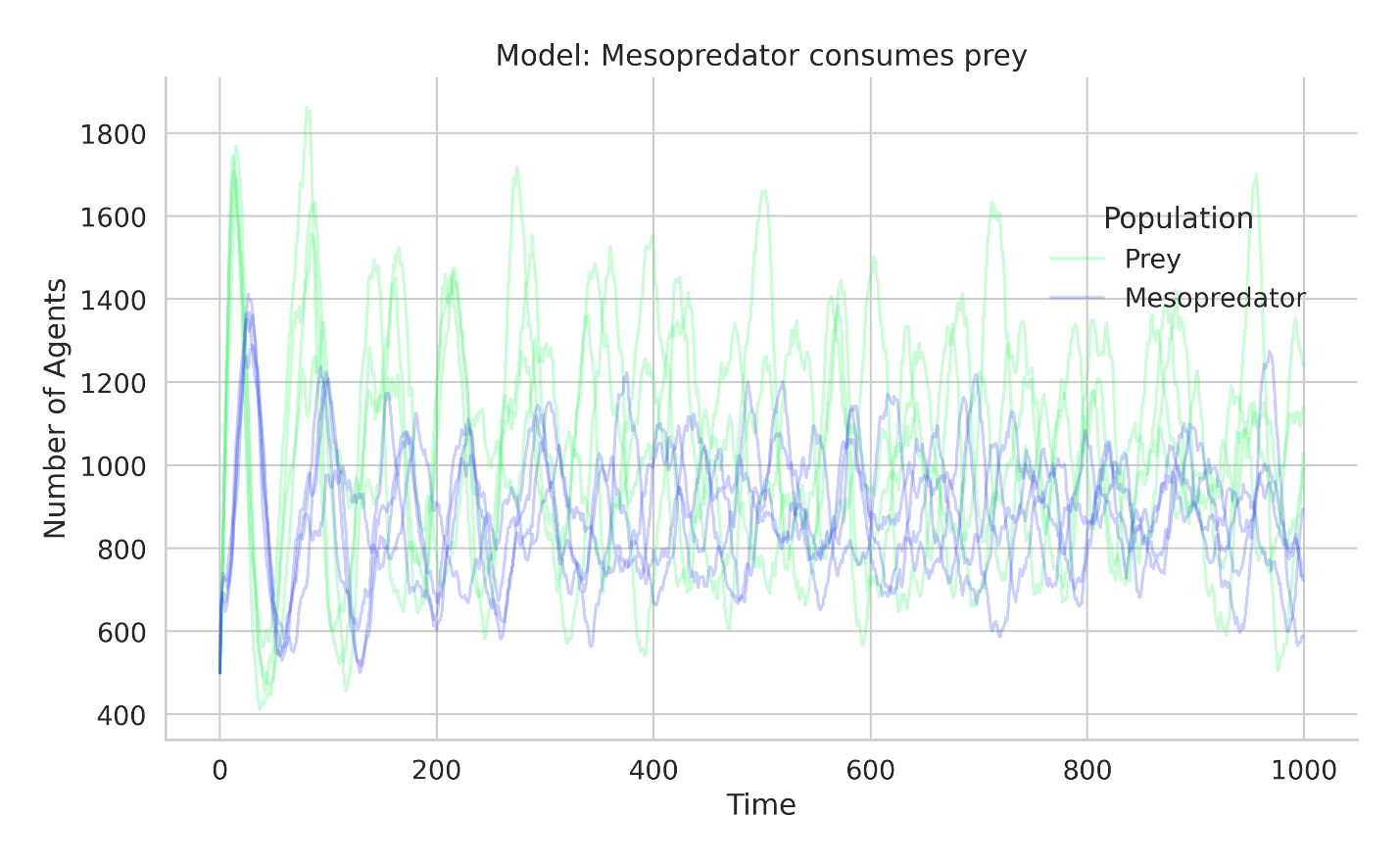

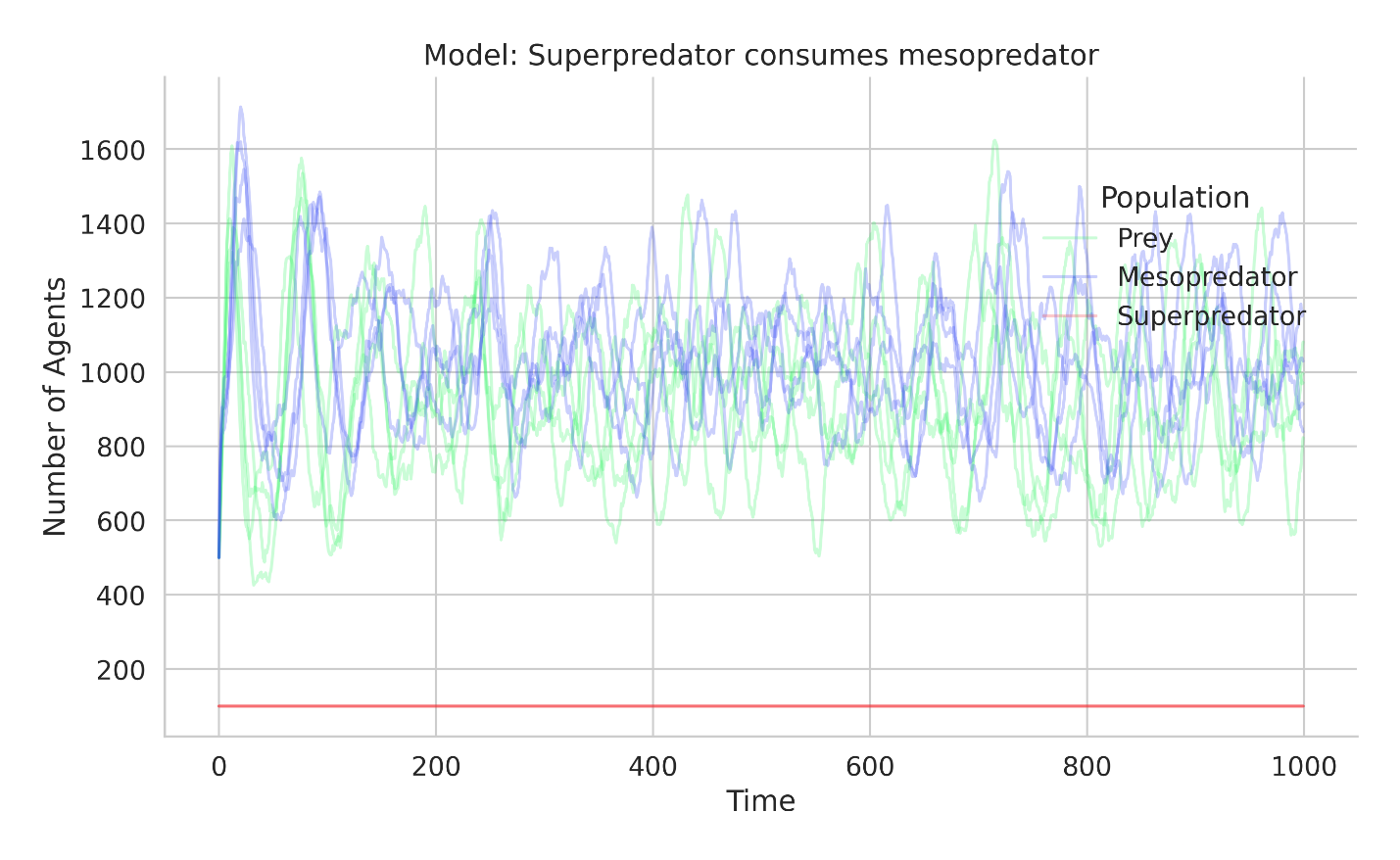

**Figure A14:** Selected examples of oscillatory population timeseries.

**Appendix B**

**Table B1:** Summary of marginal probabilities for each system state, calculated across all simulated scenarios and averaged over the complete parameter space of mesopredator and prey birth rates.

| **Scenario** | **State** | **Mean** |
| --- | --- | --- |
| Mesopredators target prey | Prey Only | 0.132 |
|  | Coexistence | 0.779 |
|  | Extinction | 0.089 |
| Apex predators target mesopredators | Prey Only | 0.140 |
|  | Coexistence | 0.812 |
|  | Extinction | 0.048 |
| Apex predators target both prey and mesopredators | Prey Only | 0.237 |
|  | Coexistence | 0.713 |
|  | Extinction | 0.050 |
| Prey respond to non-lethal superpredators | Prey Only | 0.139 |
|  | Coexistence | 0.812 |
|  | Extinction | 0.048 |
| Mesopredators respond to non-lethal superpredators | Prey Only | 0.140 |
|  | Coexistence | 0.812 |
|  | Extinction | 0.048 |
| Prey and Mesopredators respond to non-lethal superpredators | Prey Only | 0.139 |
|  | Coexistence | 0.813 |
|  | Extinction | 0.048 |
| Superperedators target prey | Prey Only | 0.131 |
|  | Coexistence | 0.471 |
|  | Extinction | 0.398 |
| Superpredators target mesopredators | Prey Only | 0.180 |
|  | Coexistence | 0.771 |
|  | Extinction | 0.048 |
| Superpredators target prey and mesopredators | Prey Only | 0.181 |
|  | Coexistence | 0.748 |
|  | Extinction | 0.071 |

**Table B2:** Summary of necessary conditions in terms of predator and prey birth rates for each possible model state by scenario.

| **Scenario** | **Variable** | **State** | **Mean** | **5th Percentile** | **95th Percentile** |
| --- | --- | --- | --- | --- | --- |
| Mesopredators target prey | Predator Birth Rate | Prey Only | 0.294 | 0.100 | 0.908 |
|  |  | Coexistence | 0.569 | 0.192 | 0.945 |
|  |  | Extinction | 0.787 | 0.265 | 1.000 |
|  | Prey Birth Rate | Prey Only | 0.316 | 0.118 | 0.835 |
|  |  | Coexistence | 0.564 | 0.192 | 0.963 |
|  |  | Extinction | 0.691 | 0.100 | 1.000 |
| Apex predators target mesopredators | Predator Birth Rate | Prey Only | 0.295 | 0.100 | 0.871 |
|  |  | Coexistence | 0.574 | 0.192 | 0.963 |
|  |  | Extinction | 0.575 | 0.137 | 0.982 |
|  | Prey Birth Rate | Prey Only | 0.326 | 0.100 | 0.871 |
|  |  | Coexistence | 0.587 | 0.210 | 0.963 |
|  |  | Extinction | 0.242 | 0.100 | 0.871 |
| Apex predators target both prey and mesopredators | Predator Birth Rate | Prey Only | 0.272 | 0.100 | 0.780 |
|  |  | Coexistence | 0.644 | 0.284 | 0.982 |
|  |  | Extinction | 0.559 | 0.137 | 0.963 |
|  | Prey Birth Rate | Prey Only | 0.526 | 0.118 | 0.982 |
|  |  | Coexistence | 0.571 | 0.192 | 0.945 |
|  |  | Extinction | 0.183 | 0.100 | 0.743 |
| Prey respond to non-lethal superpredators | Predator Birth Rate | Prey Only | 0.278 | 0.100 | 0.835 |
|  |  | Coexistence | 0.576 | 0.173 | 0.963 |
|  |  | Extinction | 0.587 | 0.155 | 0.982 |
|  | Prey Birth Rate | Prey Only | 0.336 | 0.100 | 0.871 |
|  |  | Coexistence | 0.583 | 0.192 | 0.963 |
|  |  | Extinction | 0.241 | 0.100 | 0.871 |
| Mesopredators respond to non-lethal superpredators | Predator Birth Rate | Prey Only | 0.282 | 0.100 | 0.871 |
|  |  | Coexistence | 0.578 | 0.210 | 0.982 |
|  |  | Extinction | 0.605 | 0.137 | 0.982 |
|  | Prey Birth Rate | Prey Only | 0.327 | 0.100 | 0.853 |
|  |  | Coexistence | 0.586 | 0.192 | 0.963 |
|  |  | Extinction | 0.260 | 0.100 | 0.890 |
| Prey and Mesopredators respond to non-lethal superpredators | Predator Birth Rate | Prey Only | 0.277 | 0.100 | 0.853 |
|  |  | Coexistence | 0.601 | 0.210 | 0.982 |
|  |  | Extinction | 0.578 | 0.137 | 0.982 |
|  | Prey Birth Rate | Prey Only | 0.340 | 0.100 | 0.890 |
|  |  | Coexistence | 0.584 | 0.192 | 0.982 |
|  |  | Extinction | 0.247 | 0.100 | 0.871 |
| Superperedators target prey | Predator Birth Rate | Prey Only | 0.317 | 0.100 | 0.743 |
|  |  | Coexistence | 0.401 | 0.155 | 0.651 |
|  |  | Extinction | 0.801 | 0.522 | 0.982 |
|  | Prey Birth Rate | Prey Only | 0.318 | 0.137 | 0.816 |
|  |  | Coexistence | 0.589 | 0.247 | 0.945 |
|  |  | Extinction | 0.550 | 0.118 | 0.963 |
| Superpredators target mesopredators | Predator Birth Rate | Prey Only | 0.293 | 0.100 | 0.835 |
|  |  | Coexistence | 0.592 | 0.210 | 0.963 |
|  |  | Extinction | 0.555 | 0.137 | 0.963 |
|  | Prey Birth Rate | Prey Only | 0.354 | 0.100 | 0.871 |
|  |  | Coexistence | 0.591 | 0.210 | 0.963 |
|  |  | Extinction | 0.208 | 0.100 | 0.816 |
| Superpredators target prey and mesopredators | Predator Birth Rate | Prey Only | 0.317 | 0.100 | 0.853 |
|  |  | Coexistence | 0.598 | 0.210 | 0.963 |
|  |  | Extinction | 0.554 | 0.137 | 0.963 |
|  | Prey Birth Rate | Prey Only | 0.358 | 0.137 | 0.908 |
|  |  | Coexistence | 0.607 | 0.229 | 0.963 |
|  |  | Extinction | 0.145 | 0.100 | 0.431 |

**Table B3:** Summary of dominant period of oscillation of mesopredator and prey populations under the coexistence state across scenarios.

| **Scenario** | **Population** | **Mean Period** | **SD Period** |
| --- | --- | --- | --- |
| Mesopredators target prey | Predator | 100.465 | 113.054 |
|  | Prey | 106.683 | 117.091 |
| Apex predators target mesopredators | Predator | 102.990 | 117.804 |
|  | Prey | 109.377 | 122.722 |
| Apex predators target both prey and mesopredators | Predator | 107.807 | 130.114 |
|  | Prey | 119.275 | 136.508 |
| Prey respond to non-lethal superpredators | Predator | 103.791 | 119.537 |
|  | Prey | 110.975 | 125.599 |
| Mesopredators respond to non-lethal superpredators | Predator | 103.703 | 117.902 |
|  | Prey | 110.255 | 123.996 |
| Prey and Mesopredators respond to non-lethal superpredators | Predator | 103.431 | 118.063 |
|  | Prey | 109.255 | 122.522 |
| Superperedators target prey | Predator | 116.760 | 136.532 |
|  | Prey | 130.534 | 145.726 |
| Superpredators target mesopredators | Predator | 103.902 | 130.984 |
|  | Prey | 113.214 | 136.239 |
| Superpredators target prey and mesopredators | Predator | 113.842 | 137.758 |
|  | Prey | 127.997 | 146.233 |

**Table B4**: Change in probability of possible states in the model and change in the period of population oscillations for scenarios described in table 1 of main text relative to the baseline scenario when mesopredators and prey coexist.

| State probabilities | | | | | |
| --- | --- | --- | --- | --- | --- |
| Scenario | State | Mean Effect | 5.00% | 95.00% | P(Effect > 0) |
| Apex predator consumes mesopredator | Prey Only | 0.007 | 0.005 | 0.009 | 1 |
|  | Coexistence | 0.034 | 0.031 | 0.036 | 1 |
|  | Extinction | -0.041 | -0.043 | -0.039 | 0 |
| Apex predator consumes both prey and mesopredator | Prey Only | 0.105 | 0.103 | 0.107 | 1 |
|  | Coexistence | -0.066 | -0.068 | -0.063 | 0 |
|  | Extinction | -0.039 | -0.041 | -0.037 | 0 |
| Superpredator consumes prey | Prey Only | -0.002 | -0.004 | 0.000 | 0.083 |
|  | Coexistence | -0.307 | -0.310 | -0.305 | 0 |
|  | Extinction | 0.309 | 0.307 | 0.311 | 1 |
| Superpredator consumes mesopredator | Prey Only | 0.048 | 0.046 | 0.050 | 1 |
|  | Coexistence | -0.007 | -0.010 | -0.005 | 0 |
|  | Extinction | -0.041 | -0.043 | -0.039 | 0 |
| Superpredator consumes both prey and mesopredator | Prey Only | 0.049 | 0.048 | 0.051 | 1 |
|  | Coexistence | -0.031 | -0.033 | -0.028 | 0 |
|  | Extinction | -0.019 | -0.021 | -0.017 | 0 |
| Prey respond to non-lethal superpredator | Prey Only | 0.007 | 0.005 | 0.009 | 1 |
|  | Coexistence | 0.034 | 0.031 | 0.036 | 1 |
|  | Extinction | -0.041 | -0.043 | -0.039 | 0 |
| Mesopredator respond to non-lethal superpredator | Prey Only | 0.008 | 0.006 | 0.010 | 1 |
|  | Coexistence | 0.033 | 0.031 | 0.036 | 1 |
|  | Extinction | -0.041 | -0.043 | -0.039 | 0 |
| Both prey and mesopredator respond to non-lethal superpredator | Prey Only | 0.007 | 0.005 | 0.009 | 1 |
|  | Coexistence | 0.034 | 0.032 | 0.036 | 1 |
|  | Extinction | -0.041 | -0.043 | -0.039 | 0 |
| Period of coexisting population oscillations | | | | | |
| Scenario | Agent | Mean Effect | 5.00% | 95.00% | P(Effect > 0) |
| Apex predator consumes mesopredator | Prey | 4.262 | 0.228 | 7.473 | 1 |
|  | Predator | 3.571 | 0.527 | 7.574 | 1 |
| Apex predator consumes both prey and mesopredator | Prey | 34.863 | 31.907 | 38.814 | 1 |
|  | Predator | 4.359 | 1.592 | 7.498 | 1 |
| Superpredator consumes prey | Prey | -9.346 | -12.190 | -5.718 | 0 |
|  | Predator | -7.773 | -11.424 | -2.307 | 0 |
| Superpredator consumes mesopredator | Prey | 9.732 | 7.212 | 13.519 | 1 |
|  | Predator | 1.417 | -1.648 | 5.210 | 0.76 |
| Superpredator consumes both prey and mesopredator | Prey | 36.002 | 32.544 | 38.876 | 1 |
|  | Predator | 7.088 | 3.964 | 10.467 | 1 |
| Prey respond to non-lethal superpredator | Prey | 5.264 | 1.279 | 9.615 | 1 |
|  | Predator | 3.752 | 1.248 | 7.151 | 1 |
| Mesopredator respond to non-lethal superpredator | Prey | 4.853 | 0.835 | 8.493 | 1 |
|  | Predator | 4.018 | 0.626 | 6.404 | 0.96 |
| Both prey and mesopredator respond to non-lethal superpredator | Prey | 4.186 | 0.612 | 6.297 | 0.96 |
|  | Predator | 3.751 | 0.376 | 6.133 | 1 |

**Appendix C**

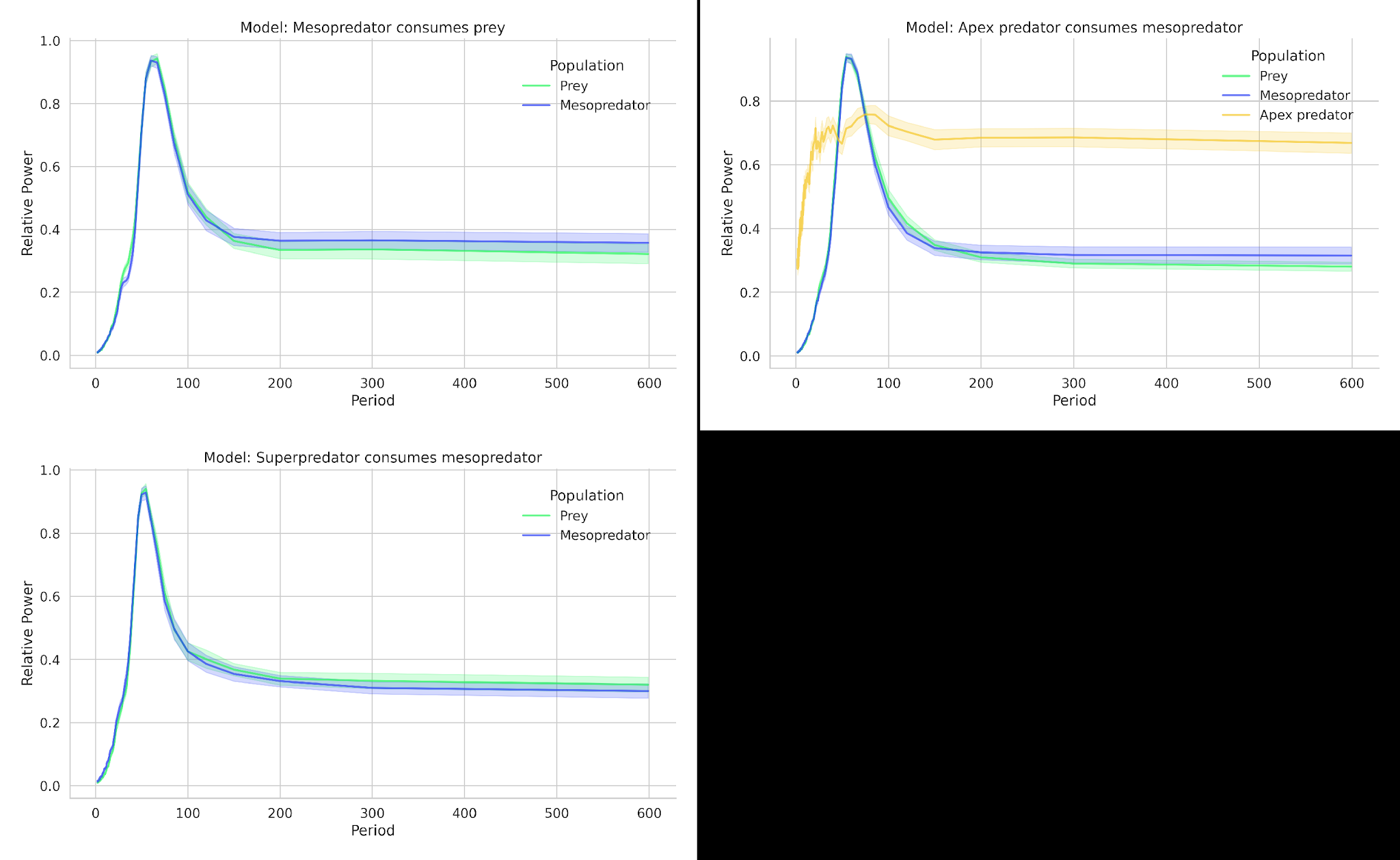

**Figure C1:** Comparing average power spectra of mesopredator and prey populations when *bprey* *= 0.56* and *bmesopredator = 0.2* between baseline scenario and when superpredators kill mesopredators.

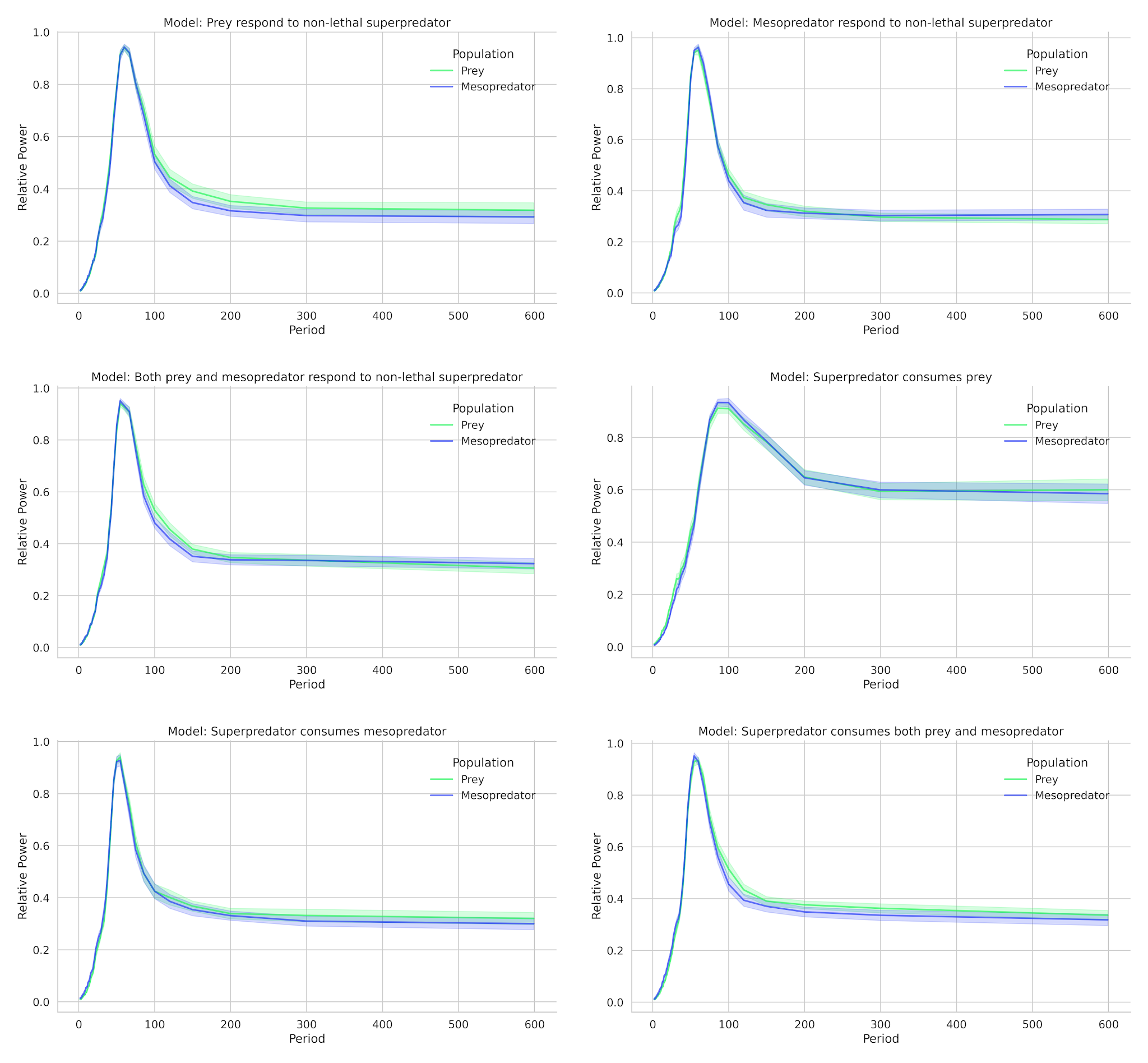

**Figure C2:** Comparing average power spectra of mesopredator and prey populations when *bprey* *= 0.56* and *bmesopredator = 0.2* between lethal and non-lethal superpredators targeting prey, mesopredators or both.

**Appendix D**

**Table 1**: Summary of past models of multi-trophic risk in predator – prey systems.

| **Authors and year** | **Type** | **Description** |
| --- | --- | --- |
| Wang et al., 2016 | Continuous mean – field model | Lotka – Volterra based model with cost of fear incorporated into prey production. |
| Mishra et al. 2021; Verma et al. 2021; Pal et al, 2019; Wang et al. 2017 | Continuous mean – field model | Modifications of Wang et al.’s (2016) with various predator response functions |
| Mitchell and Lima, 2002 | Agent based model | Spatially explicit model incorporating vigilance and movement strategies to mitigate predation risk. |
| Cain and Mitchell, 2021; Luttbeg and Schmitz, 2000; Mitchell, 2018 | Agent based/Game theoretic model | 2 patch game between predator and prey. |
| Brown et al. 1999 | Optimal foraging model | Patch based foraging model where prey optimise energy gain and minimise predation risk. |
| Dong and Li, 2021; Duan et al. 2019; Upadhyay and Mishra, 2019; Souna et all. 2021 | Reaction – diffusion model | Model of diffusive predators, prey or both with fear incorporated as in Wang et al. (2016) |
